# Peer pressure from a *Proteus mirabilis* self-recognition system controls participation in cooperative swarm motility

**DOI:** 10.1101/490771

**Authors:** Murray J. Tipping, Karine A. Gibbs

## Abstract

Colonies of the opportunistic pathogen *Proteus mirabilis* can distinguish self from non-self: in swarming colonies of two different strains, one strain excludes the other from the expanding colony edge. Predominant models characterize bacterial kin discrimination as immediate antagonism towards non-kin cells, typically through delivery of toxin effector molecules from one cell into its neighbor. Upon effector delivery, receiving cells must either neutralize it by presenting a cognate anti-toxin, as would a clonal sibling, or suffer cell death or irreversible growth inhibition, as would a non-kin cell. Here we expand this paradigm to explain the non-lethal Ids self-recognition system, which stops access to a cooperative social behavior in *P. mirabilis* through a distinct mechanism: selectively and transiently inducing nonself cells into a lifestyle incompatible with cooperative swarming. This state is characterized by reduced expression of genes associated with protein synthesis, virulence, and motility, and also causes non-self cells to tolerate previously lethal concentrations of antibiotics. We found that entry into this state requires a temporary activation of the stringent response in non-self cells and results in the iterative exclusion of non-self cells as a swarm colony migrates outwards. These data clarify the intricate connection between non-lethal recognition and the lifecycle of *P. mirabilis* swarm colonies.

## Introduction

Organisms rarely live in complete isolation. Living in a community can provide benefits to each individual. However, there is a constant balance between the interests of individuals and the maintenance of community-wide advantages. A stable evolutionary strategy is for individuals to preferentially direct advantages to close kin (1–3). This behavior, known as kin discrimination, has been the subject of focused study.

Several examples of kin discrimination in bacteria have been elegantly described, including those mediated by Type IV (4), Type VI (5,6) and Type VII (7) secretion system based effector exchange, contact-dependent inhibition (CDI) (8, 9), and the MafB toxins of the *Neisseria* (10). One common thread between these systems is that they characterize discrimination as immediate and irreversible antagonism towards cells or strains that are non-kin, typically through delivery of lethal toxin effector molecules. Upon effector delivery, receiving cells must either neutralize it by presenting a cognate anti-toxin or suffer immediate negative consequences, typically cell death (5) or permanent inhibition of growth (8). Here we describe an expansion of these mechanisms: the Ids self-recognition system mediates kin discrimination in *Proteus mirabilis* by selectively inducing non-self cells into a lifestyle incompatible with swarming, thereby controlling access to a cooperative social behavior.

*P. mirabilis*, a major cause of recurrent complicated urinary tract infections (11), engages in several sophisticated social behaviors such as swarming on rigid surfaces. Swarms are formed by many elongated (approximately 10 - 80 μm) “swarmer” cells moving cooperatively, allowing for colony expansion over centimeter-scale distances. Rounds of swarming are interspersed with periods of non-expansion termed “consolidation”. The oscillation between swarming and consolidation leads to a characteristic pattern of concentric rings on higher percentage agar plates (12). Effective *P. mirabilis* swarming relies on the ability of swarmer cells to form large rafts that together move more quickly than isolated individuals (13). Rafts are fluid, transient collectives that cells frequently enter and exit. As such, an individual cell interacts with many different neighbors through the lifetime of a swarm. During swarming, *P. mirabilis* cells can communicate with each other by exchanging proteins through contact-dependent secretion systems (14, 15). These signals in turn cause emergent changes in swarm behavior (16, 17).

*P. mirabilis* swarms exhibit the ability to recognize self in several ways. The oldest known example is Dienes line formation: two swarms of the same strain merge into a single swarm upon meeting, while two swarms of different strains instead form a human-visible boundary (18, 19). More recently, the phenomenon of territorial exclusion was described: in a mixed swarm comprised of two different strains, one strain is prevented from swarming outwards by the other (14). Clonal swarms of *P. mirabilis* have a coherent self identity, minimally mediated by the Ids system encoded by six genes, *idsA-F*. Deletion of the *ids* locus results in the mutant strain no longer recognizing its wild-type parent as self (20). Many of the molecular mechanisms governing Ids-mediated self recognition have been described in detail (Figure 1A). However, how the Ids system functions in local behaviors has remained elusive.

Briefly, two proteins, IdsD and IdsE, govern self identity. IdsD is transferred between cells in a Type VI secretion system (T6SS)-dependent fashion; disruption of the T6SS prevents all Ids signal transfer (17, 21). A cell in a swarm is considered to be self if it produces an IdsE protein that can bind IdsD proteins sent from neighboring cells. Disruption of these IdsD-IdsE interactions, either through deletion of *idsE* or through swapping *idsD* or *idsE* with variants from another *P. mirabilis* strain, result in strains that display extreme attenuation in swarm expansion without loss of viability (16,17). The IdsD protein must be incoming; endogenous IdsD and IdsE proteins produced within a single cell do not impact self-recognition behaviors (21). Conditions that lead to non-self recognition are described as “Ids mismatch”.

Crucially, territorial exclusion by Ids does not affect viability. Excluded cells have been shown to grow and divide at a rate comparable to non-excluded cells (17). Yet Ids has been described as a toxin-antitoxin system (22, 23). This characterization is inconsistent with experimental data and is likely due to the reliance on the T6SS for transport of IdsD as the T6SS is often characterized as a lethal toxin delivery mechanism (24). A mechanistic description of Ids-mediated recognition is needed to reconcile the data and would also provide a model for other non-lethal mechanisms that might be attuned for surface-dwelling swarm migration.

Here we show that even though Ids recognition signal transfer happens while cells are actively migrating as a swarm, the recognition response is delayed until cells have stopped moving. We show that recognition of non-self is at least partially mediated by ppGpp levels within the cell and this contributes to a concerted shift of cells into a distinct, antibiotic-tolerant state that is incompatible with continuation of swarming. The increased antibiotics tolerance is due to the cell-to-cell transfer of IdsD. We found that this Ids-induced state is short-acting; induction requires continuous cell-cell interactions. In the context of a swarm, the collective consequence is an iterative winnowing of the non-self cells from the swarm fronts during periods of no active migration. We posit that the cell-cell communication of these non-lethal factors acts as a control system during swarm expansion by diverting non-self from developing into swarm-compatible cells and thus preventing non-self cells from taking part in cooperative swarm behavior.

## Results

### Ids mismatch during swarming induces a distinct transcriptional state

We initially sought to determine the method through which Ids caused non-self cells to be territorially excluded from swarms. Given the lack of lethality and the stark attenuation of swarm colony expansion observed during Ids-mediated territorial exclusion (14, 17), we hypothesized that an Ids mismatch caused broad changes in gene expression of the recipient cell. Ids mismatch is defined here as transcellular communication of IdsD to a recipient cell lacking a cognate IdsE (Figure 1A). Therefore, we performed RNA-Seq differential expression analysis using conditions that would produce either self or non-self interactions, all within genetically equivalent backgrounds. We isolated total RNA from cells undergoing consolidation, because consolidation is the swarm development stage most tightly connected to major transcriptional changes (25).

As a baseline, we first compared transcriptional profiles between cells from clonal swarms of wildtype and independently, of a derived mutant strain lacking the *ids* genes (*BB2000::idsΩCm* (20), herein referred to as “Δ*ids*”). The swarm colonies of both strains expand equivalently and have no notable morphological differences (20). 10 genes showed significant (fold-change > log_2_ 1.5, p < 0.05) differential regulation in clonal swarms of the Δ*ids* strain as compared to wildtype, six of which were the Δ*ids* genes deleted in the construction of the Δ*ids* strain (Figure 1B). A complete list of differentially regulated genes is found in Supplementary Table 1. We concluded that there is little change in gene expression between wildtype and the Δ*ids* strain.

We next considered differences in strains experiencing Ids mismatch. We examined three different conditions: a clonal swarm in which every cell lacked the *idsE* gene (CCS06), a clonal swarm in which every cell contained an IdsE protein unable to bind transferred IdsD proteins (CCS02), and cells of a Δ*ids*-derived strain constitutively producing Green Fluorescent Protein GFPmut2 (Δ*ids-GFP*) that were isolated from a 1:1 co-swarm with wildtype through fluorescent-activated cell sorting (FACS), henceforth termed “co-swarmed Δ*ids*”. All strains have previously been verified and characterized (17, 20, 21). As the two clonal swarm colonies have attenuated expansion, we were only able to harvest whole colonies as visible consolidation phases were less distinct. RNA-Seq differential expression analysis was performed on cells from each condition as compared to the appropriate control samples: clonal Δ*ids* for co-swarmed Δ*ids* or a clonal Δ*ids*-derived swarm in which every cell expressed *in trans* plasmid-encoded *ids* genes (CCS01, Δ*ids*_pidsBB_) for the clonal swarms. Broad changes to relative transcript abundances were apparent for each condition: 231 genes in CCS06, 457 genes in CCS02, and 836 genes in co-swarmed Δ*ids*, which represents approximately 6%, 13%, and 23% of total genes, respectively (Figure 1, Supplementary Tables 2, 3 and 4).

Trends were apparent across the datasets. We observed a concerted decrease in transcripts for class I, II, and III genes for flagellar synthesis such as *flhDC*, *filA*, and *fliC*. We also observed a decrease in transcripts for many genes associated with protein synthesis, such as the 50S ribosomal protein *rplT* and 30S ribosomal protein *rpsP*, along with ribosomal-associated elongation factors such as EF-Tu. Several genes involved in respiration, including those of the F_0_F_1_-ATP synthase, also had significantly fewer transcripts as did those transcripts for different virulence-associated protein families (26–31), such as *hpmA*, *umoA*, and *zapD*. Overall, fewer genes had an increased relative abundance as compared to the control strains: 109 genes in CCS06, 56 genes in CCS02, and 332 genes in co-swarmed Δ*ids*, respectively. Within these genes, several endogenous toxin-antitoxin systems displayed an increased relative abundances of tran-scripts, including genes that encode homologous proteins to *Escherichia coli* YfiA (also known as RaiA) (32) and to toxin/antitoxin pairs ParE/CC2985 (33) and Phd/Doc (34). There was also an increased relative abundance for several fimbriae families, including genes encoding MR/P and P-like fimbria. A large proportion of differentially regulated genes were proteins of unknown function.

**Fig. 1.**
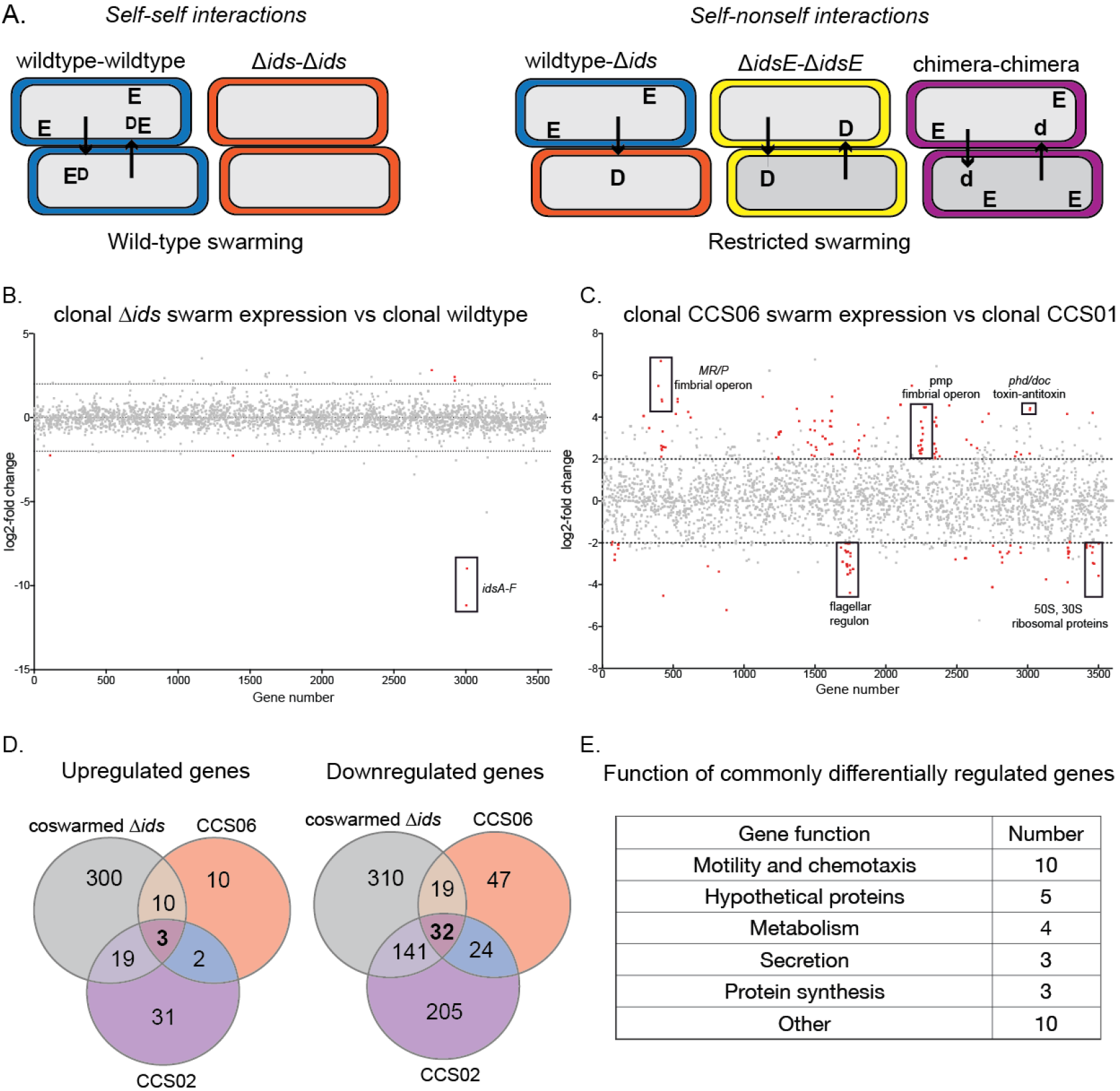
Ids mismatch during swarming causes large transcriptional shifts. (A) Cartoons show self versus non-self interactions as defined by the Ids state in and between cells. The post-transferred state of IdsD is depicted. (B) A graphical representation of genes differentially regulated in consolidator cells isolated from clonal Δ*ids* swarms versus consolidator cells from clonal wild-type swarms. The X-axis corresponds to each gene on the *P. mirabilis* BB2000 chromosome. The Y-axis corresponds to log_2_-fold difference in relative transcript abundance. Significantly differentially regulated genes, defined as fold-change > log_2_ 1.5, p < 0.05, are labeled in red. All data are included. The genes *idsA-F*, which were deleted to construct the Δ*ids* strain, are labeled. (C) A graphical representation of genes differentially regulated in whole swarms of strain CCS06 versus consolidators from strain CCS01, constructed similarly to that of (A). Labeled genes are discussed in the text. (D) Venn diagrams showing genes similarly differentially regulated in three separate swarm conditions for an Ids mismatch between cells: co-swarmed Δ*ids*, clonal CCS06, and clonal CCS02. Two biological repeats were performed. (E) Annotated (taken from KEGG and COGG databases) functions of the 35 genes for which relative transcript abundance was significantly different from that of a clonal wild-type population (C): co-swarmed Δ*ids*, clonal CCS06 and clonal CCS02.

A subset of differentially abundant genes was shared among all three datasets; these were representative of families with a decreased relative abundance of transcripts in each Ids mismatch strain. Three genes had increased relative abundance in transcripts as compared to wildtype; 32 genes had decreased (Figure 1D). The 32 genes with decreased relative transcripts included those involved in motility, chemotaxis, ribosomal proteins, and metabolism (Figure 1E). Of the three genes with increased relative transcripts, two encode the Phd/Doc endogenous toxin-antitoxin system. The third gene is *rob*, which has been associated with the induction of low-metabolism states in other bacterial species (35). Thus, cells under the influence of incoming IdsD, and without a cognate IdsE protein present, enter a state distinct from either wildtype or the Δ*ids* cells participating in a normal swarm cycle. Transcriptional shifts of these types are often associated with entry into altered states (e.g., increased antibiotics tolerance) that are induced by a variety of environmental and temporal cues, including nutrient and membrane stress (36).

### Cells experiencing Ids mismatch display increased tolerance to lethal concentrations of antibiotics

We hypothesized that the changes in Ids-excluded cells might also result in the secondary effect of increased an-tibiotic tolerance. We conducted antibiotic tolerance assays on wildtype, the Δ*ids* strain, and a third strain containing an in-frame deletion of the chromosome-encoded *idsE* within the wildtype background (21). This chromosomal *idsE* deletion mutant strain (Δ*idsE*) was used to avoid confounding factors associated with antibiotics and plasmid maintenance. We predicted that swarms of the Δ*idsE* strain would display increased antibiotics tolerance as compared to wildtype and the Δ*ids* strain.

**Fig. 2.**
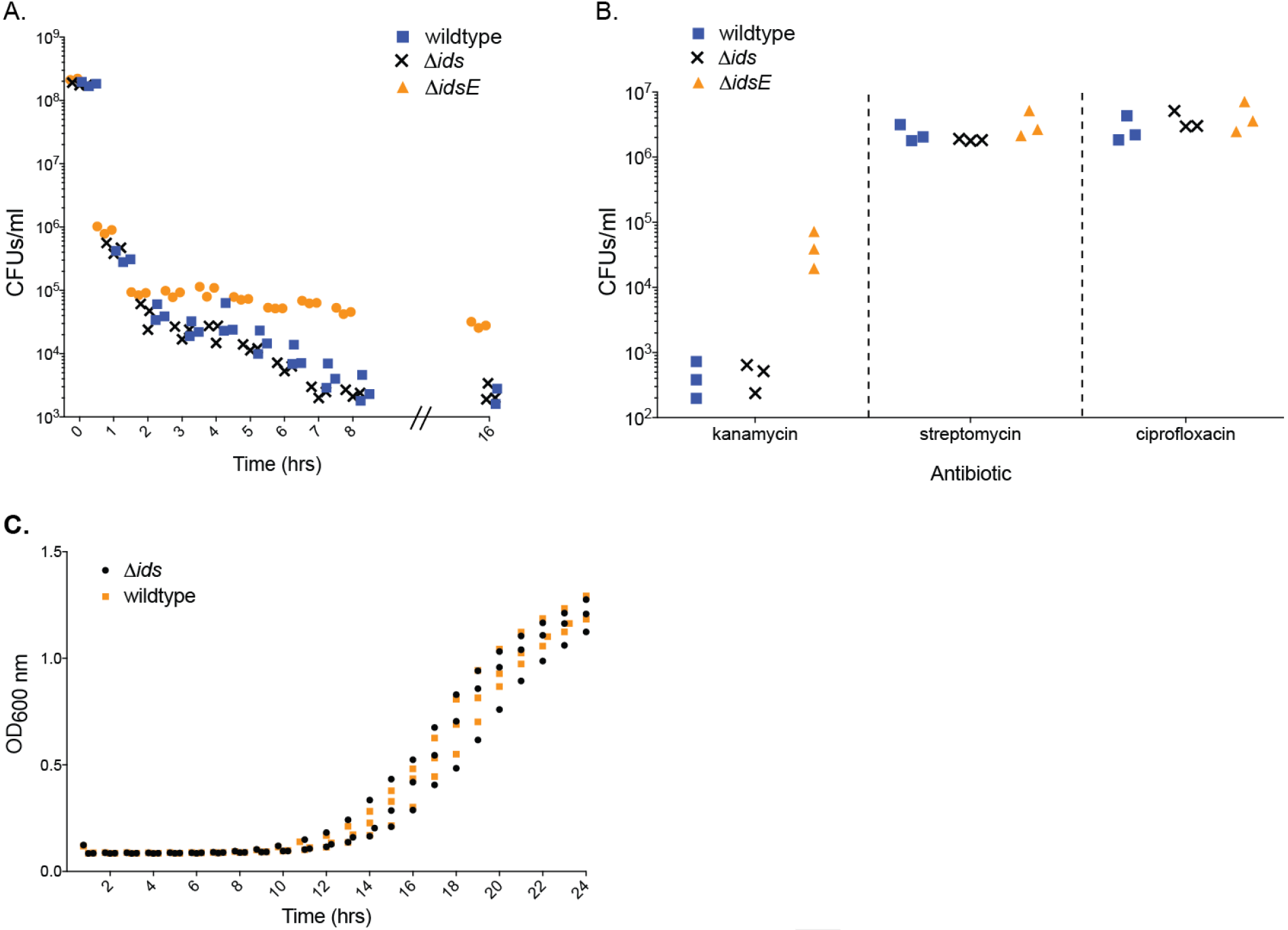
Cells experiencing Ids mismatch display transient tolerance to lethal concentrations of antibiotics. (A) Killing curves for cells of Δ*ids* (black “x”), Δ*idsE* (orange triangle) and wildtype (blue square) harvested from swarm plates and exposed to 100 μg ml^-1^ ampicillin. Note that the Y-axis is a logarithmic scale. (B) Survival of strainsΔ*ids* (black “x”), Δ*idsE* (orange triangle) and wildtype (blue square) after swarm colonies were harvested and exposed to 60 μg ml^-1^1 kanamycin (left), 50 μg ml^-1^ streptomycin (middle), and 1 μg ml^-1^ ciprofloxacin (right) for 12 hours. (C) Territorial exclusion does not result in long-term growth defects. Optical density at 600 nm (OD_600_) was measured over time for liquid cultures of wildtype and the co-swarmed Δ*ids* strain. Liquid cultures were inoculated usi g cells isolated by FACS from co-swarm colonies where the Δ*ids* strain had been actively excluded from the swarm front.

For the assays, we harvested cells from independent, clonal swarms of either wildtype, the Δ*ids* strain, or the Δ*idsE* strain after the entry to the third swarm ring. Cells were resuspended in LB media and immediately subjected to 100 μg ml^-1^ ampicillin exposure. Samples were extracted for viability assays on fresh media lacking antibiotics (Figure 2A). No clear difference was observed between wild-type and the Δ*ids* strain at any timepoint. Wildtype and the Δ*ids* strain exhibited a killing of approximately 10^5^-fold after eight hours, while the Δ*idsE* strain experienced a killing of approximately 10^4^-fold (Figure 2A). Under these conditions, the Δ*idsE* strains had an approximately 50-fold increase in survival as compared to wildtype, even after sixteen hours incubation in ampicillin.

To investigate whether tolerance was against multiple antibiotics, we repeated this assay with three additional antibiotics: the fluoroquinolone ciprofloxacin and the aminoglycosides streptomycin and kanamycin. No difference between wildtype, the Δ*ids* strain, and the Δ*idsE* strain was observed in viable cell counts following 50 μg ml^-1^ streptomycin or 1 μg ml^-1^ ciprofloxacin exposure (Figure 2B). These antibiotics resulted in much lower rates of cell killing as compared to ampicillin, which might reflect a higher native resistance of *P. mirabilis* to these drugs (37, 38). However, the Δ*idsE* strain showed an increased number of viable cells as compared to wildtype or the Δ*ids* strain 12 hours after 60 μg ml^-1^ kanamycin incubation (Figure 2B). Therefore, the Δ*idsE* strain displays increased tolerance to both kanamycin and ampicillin under these conditions, indicating resistance to multiple, but not all, antibiotics.

Since Ids functions through cell-cell contact-dependent secretion of the identity marker IdsD (17), we tested whether IdsD secretion was required for antibiotic tolerance to emerge. We performed this assay using MJT01, which is an Δ*idsE*-derived strain containing a non-functional T6SS (17, 39) and so does not secrete IdsD. No clear difference was observed between this constructed strain and wildtype over three biological repeats (Supplementary Figure 1). To examine whether social exchange was causative, we tested cells of the Δ*idsE* strain that were grown to stationary phase with shaking in liquid. Cells secrete IdsD into growth media (14). We observed no difference in antibiotics tolerance for the Δ*idsE* strain as compared to wildtype over three biological repeats (Supplementary Figure 2). Thus, an Ids mismatch, caused by the transfer of IdsD between neighboring cells, resulted in a transcriptional shift into a non-swarming state with increased antibiotic tolerance.

**Fig. 3.**
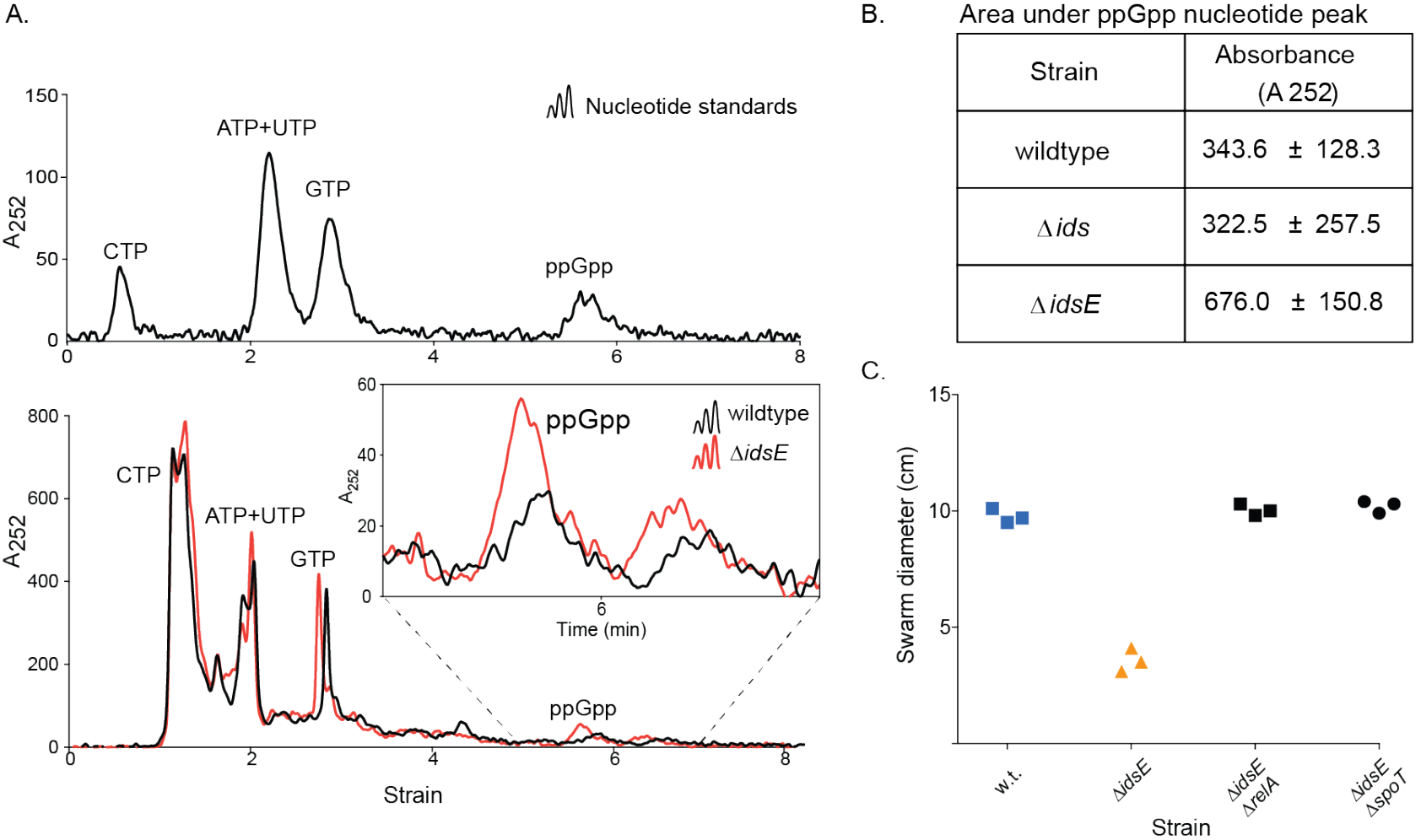
Cells affected by Ids mismatch induce an increase in ppGpp; the *relA* and *spoT* genes are necessary to induce Ids-mediated territorial exclusion. (A) Representative chromatograms showing UV absorbance (252 nm) over time of a SAX-HPLC column run. Upper graph shows peaks from 2 mmol nucleotide standards (ATP, CTP, GTP, UTP) and 0.5 mmol ppGpp. Lower graph shows peaks from running purified nucleotides from wildtype (black) and the Δ*idsE* strain (red) samples purified from swarm colonies. Inset shows the peak corresponding to absorbance from ppGpp alarmone molecule. (B) Mean ppGpp absorbances from three biological repeats of samples from wildtype, the Δ*ids* strain, and the Δ*idsE* strain, which were calculated by integrating areas under peaks. Errors show standard deviations. C. Swarm colony radius expansion of wildtype and the Δ*idsE*, Δ*idsE*Δ*relA*, and Δ*idsE*Δ*spoT* strains after 16 hours of swarming. All replicates are shown.

We considered that this Ids-mediated antibiotic tolerance might be due to entry into a persistent, irreversible dormant state. As such, we examined the dynamics of exit from Ids-induced low-swarming states by setting up co-swarms where the GFP-producing Δ*ids* strain (Δ*ids*-GFP) was inoculated with an equal amount of a wild-type strain constitutively producing DsRed (wildtype-DsRed). We let the co-swarms progress to the third swarm ring and then harvested the swarms; cells of each strain were immediately sorted by fluorescence-activated cell sorting (FACS). Equal numbers of particles of each strain were inoculated in LB media, and growth was measured for 24 hours through optical density at 600 nm. No differences between co-swarmed wildtype-DsRed and co-swarmed Δ*ids*-GFP were observed at any time-point (Figure 2C). Ids mismatch-induced effects therefore are transient outside of continual contact-mediated pressure. To confirm this, we performed the growth curve assays with the Ids mismatch strain CCS02, comparing it against strain CCS01 (Supplementary Figure 3). Again, no difference was observed. Therefore, an individual cell shifts into a distinct transcriptional state when it has received nonself signals from the surrounding cells; this cell state is temporary and reversible.

### Ids recognition mechanisms induce and require the stringent response

Entry into an antibiotic-tolerant state has been linked in other bacteria to the stringent response (36, 40), mediated by the alarmone messenger molecule (p)ppGpp. Although the stringent response has not been studied in *P. mirabilis*, the genome for wild-type BB2000 contains the two canonical genes for production and degradation of ppGpp, *relA* and *spoT* (41). Therefore, we directly measured total ppGpp quantities in cells harvested from clonal wildtype, clonal Δ*ids*, and clonal Δ*idsE* swarms experiencing Ids mismatch using a recently described high performance liquid chromatography (HPLC)-based technique (42). Nucleotide samples were purified from swarm cultures, separated by HPLC, and quantified by measuring UV absorbance spectra. Example absorption traces taken from wildtype and the Δ*idsE* samples can be seen in Figure 3A. Based on three biological repeats of ppGpp measurements (Figure 3B), the samples from the Δ*idsE* strain displayed significantly increased ppGpp levels compared to wildtype and the Δ*ids* strain.

These results were suggestive of a link between ppGpp levels and territorial exclusion. However, it did not rule out the possibility that the increased ppGpp was a consequence of other changes following transfer of unbound IdsD as opposed to a causative factor upstream of the transcriptional and physiological changes we observed. We there examined whether *relA* or *spoT* is necessary to ameliorate effects of unbound IdsD accumulation, specifically focusing on the Δ*idsE* swarm deficiency. Deletions of *relA* or *spoT* can prevent ppGpp accumulation and as such, prevent cells from activating the stringent response as discussed in (43).

**Fig. 4.**
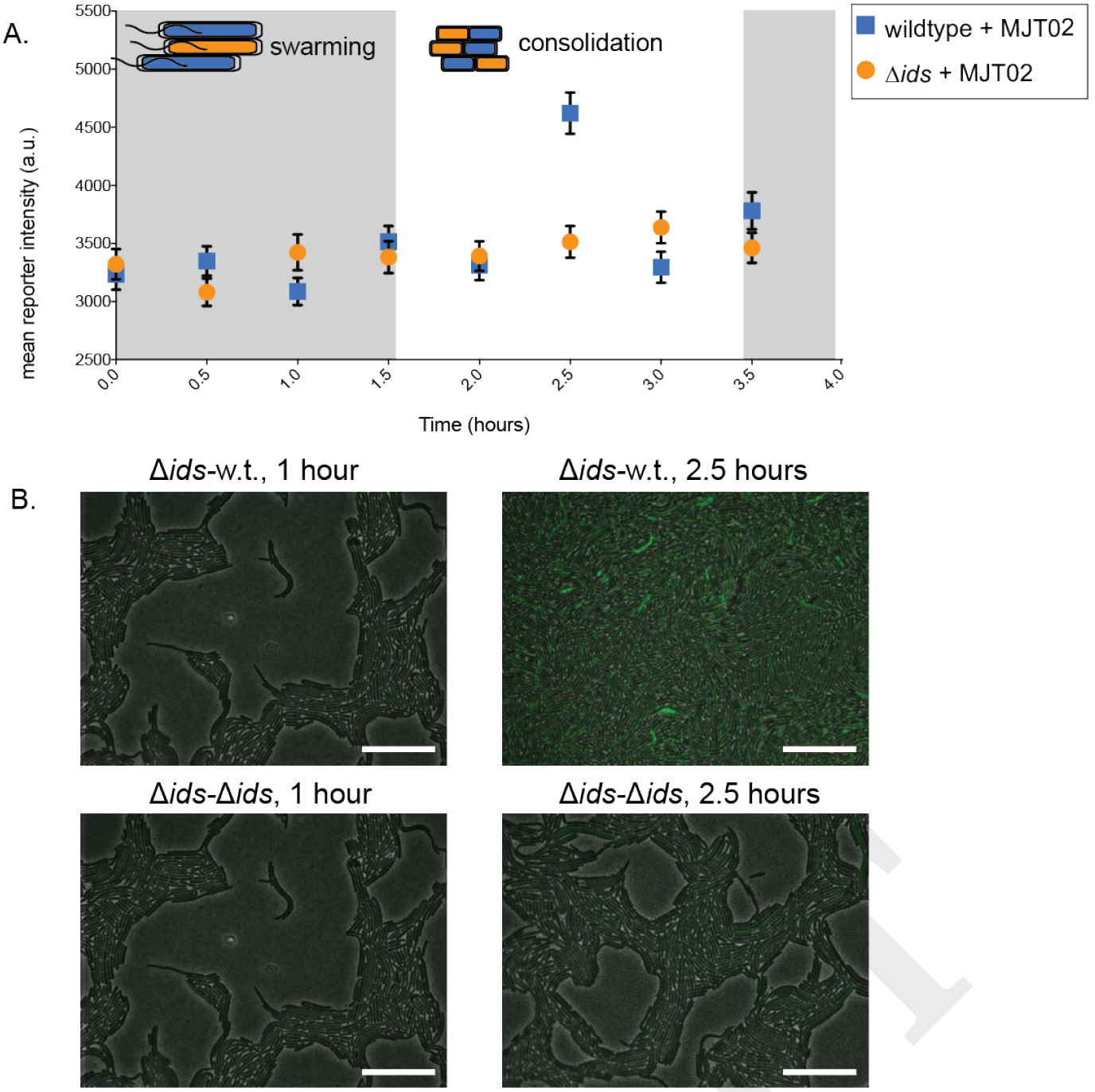
The transcriptional shift observed in cells experiencing Ids mismatch occurs during consolidation periods between swarming. (A) A time-course graph showing mean swarm fluorescence intensity overtime fortwo conditions: Δ*ids* carrying a chromosomal BB2000_0531 fluorescence reporter (MJT02, green ovals) co-swarmed with wildtype (MJT02:wildtype, blue) or co-swarmed with the Δ*ids* strain (MJT02:Δ*ids*, orange). Three biological repeats were performed. Fluorescence was measured over equivalent areas for each experimental condition. Error bars are standard deviations; a.u. means arbitrary units. (B) Representative phase contrast/Venus fluorescence overlay microscopy images (100x magnification) MJT02:wildtype (top) and MJT02:Δ*ids* (bottom) co-swarms. Images shown are of the first active swarm phase (left, 1 hour) and the first period of consolidation (right, 2.5 hours). Wildtype is abbreviated w.t. Scale bars = 10 μm.

We constructed two Δ*idsE*-derived strains with chromosomal deletions of either *relA* or *spoT*. We tested the swarm proficiency of these strains as compared to that of wildtype and the parent Δ*idsE* strain. Both newly constructed strains were equally proficient in swarming as compared to wildtype and more proficient than the parent Δ*idsE* strain (Figure 3C). Deletion of either *relA* or *spoT* in wildtype and the Δ*ids* strain did not alter swarm migration. Thus, both *relA* and *spoT* are needed for Ids mismatch to be effective.

### The distinct transcriptional shift in cells with an Ids mismatch occurs during consolidation periods

The small molecule ppGpp could allow for a rapid and transient response. Such temporal dynamics are consistent with the observation that a response to Ids mismatch is only present under consistent pressure from neighboring non-self cells (Figure 2). We reasoned then that the induction of low metabolism by Ids mismatch was either spatially or temporally limited. To interrogate this model, we took advantage of the genes newly identified as being induced in the presence of an Ids mismatch (Figure 1) to develop a fluorescent transcriptional reporter system. A gene encoding a variant of the fluorescent protein Venus (44) was engineered to be inserted immediately downstream of the gene *BB2000_0531*, resulting in Venus production being controlled by the upstream promoter. The *BB2000_0531* gene displayed increased expression under Ids mismatch conditions (Figure 1B, Supplementary Tables 2, 3, 4). This reporter construct was added to the chromosome of the Δ*ids* strain, resulting in strain MJT02; this strain had no apparent defects.

We performed fluorescence microscopy time-course experiments on mixed swarms to measure transcriptional changes associated with *BB2000_0531* over the course of a swarm-consolidation cycle. Two co-swarm conditions were used. In the first, a mixed culture of 50% MJT02 and 50% the Δ*ids* strain was used to inoculate swarm-permissive agar; in the second, a mixed culture of 50% MJT02 and 50% wildtype-DsRed was used. Venus fluorescence intensity was measured at 30-minute time-points in swarm areas, and the mean fluorescence was calculated. The fluorescence intensity for both co-swarm conditions was graphed (Figure 4A); representative images are in Figure 4B. A temporal spike in fluorescence associated with *BB2000_0531* correlated with the consolidation cycle and was only apparent when Δ*ids*-derived cells were intermingled with wild-type cells.

In addition to causing territorial exclusion in mixed swarms, Ids mismatch determines boundary formation after collision between two clonal swarms (20). Boundary formation is a complex process involving the contribution of chemical and physical factors as well as lethal and non-lethal systems (14, 15, 20, 45). To test whether equivalent transcriptional shifts were observed during the initial stages of boundary formation, when cells of each strain are in contact with one another, we measured fluorescence intensity associated with *BB2000_0531* in MJT02 swarms following collision with wildtype and Δ*ids* swarms. We observed a mean increase in fluorescence intensity over several hours after encountering wildtype, but not the Δ*ids* strain (Figure SF4). This increase in fluorescence intensity occurred before a boundary was visually apparent. In fact, formation of a visible boundary between the two strains did not occur for a further 12-18 hours after the end of this experiment, which is consistent with previous observations (18). Therefore, the Ids mismatch induces a response in the initial stages of boundary formation, once cells from different populations have interacted and before a separation is visible.

**Fig. 5.**
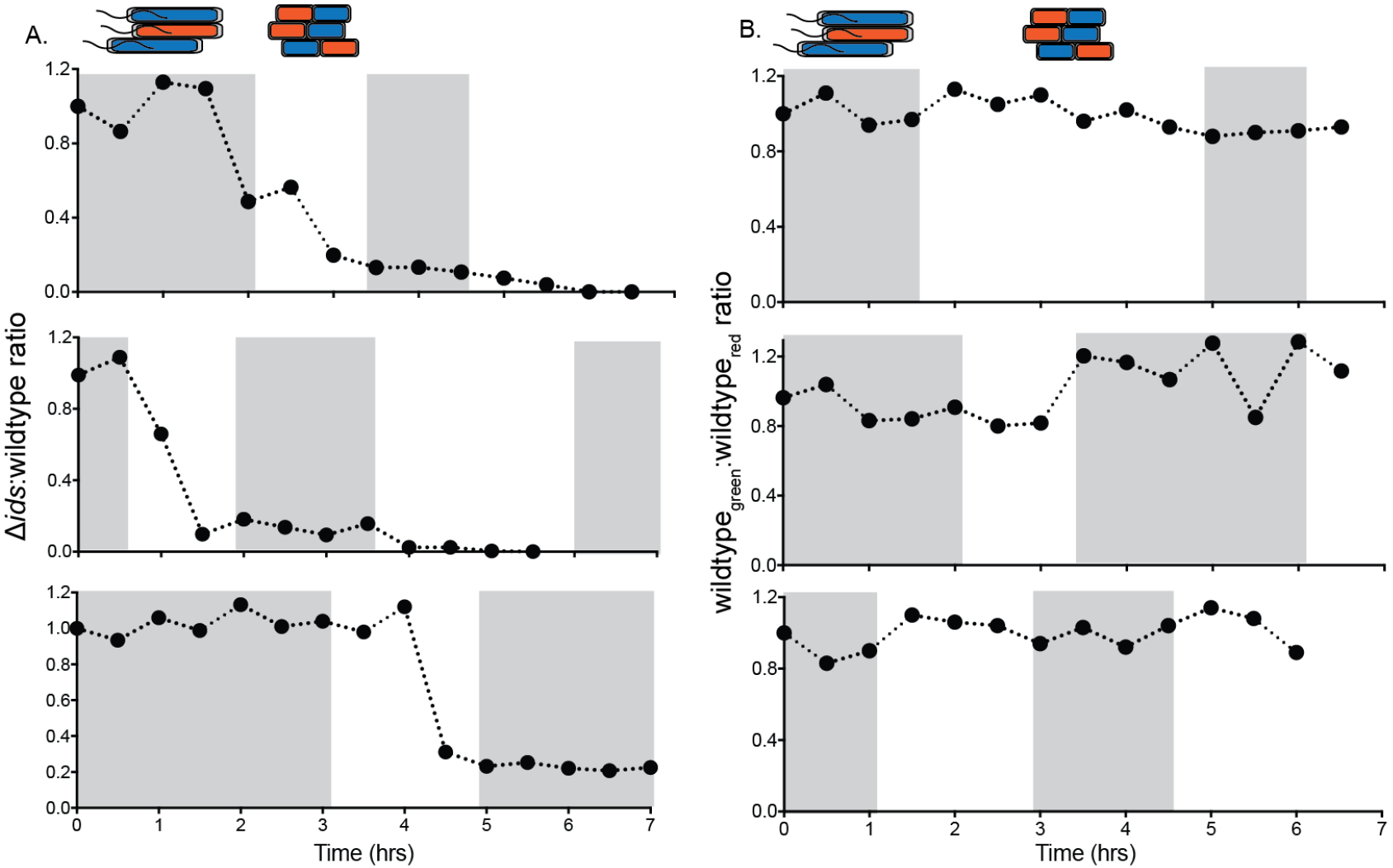
The Δ*ids* strain is excluded during consolidation between periods of swarming. 1:1 co-swarms of Δ*ids*-GFP and wildtype-DsRed (A) or wildtype-GFP and wildtype-DsRed (B) were analyzed for GFP-associated and DsRed-associated fluorescence over the course of outward migration and through progression of the swarm-consolidation cycle. Each data point indicates the proportion of GFP-producing cells measured at the swarm front at a given time after swarm emergence. Periods of active swarming were determined by eye and are highlighted in gray. Three biological repeats were performed, and each is shown.

### Non-self cells are iteratively winnowed from the swarm front

Swarming is fundamentally a collective behavior, and the spatial expansion of a wild-type swarm is connected to the oscillatory developmental cycle of outward migration and non-motile consolidation. We reasoned from these data that the Ids system likely impacts local cell-cell interactions at the boundary and within an expanding swarm. Therefore, we examined territorial exclusion *in situ* using epifluorescence microscopy and utilized co-swarms constructed with equal ratios of the Δ*ids*-GFP and wildtype-DsRed strains. The control co-swarms consisted of an equal ratio of wildtype-DsRed and wildtype constitutively producing GFPmut2 (wildtype-GFP). Once swarmer cells emerged from the inoculum, the proportion of cells expressing each fluorophore was measured at half-hour intervals. The developmental stages of active outward motility versus no outward motility (i.e., consolidation) were noted by eye.

We calculated the fluorophore ratios over time for both the Δ*ids*-GFP:wildtype-DsRed and wildtype-GFP:wildtype-DsRed co-swarms for three biological repeats (Figure 5). The GFP/DsRed ratios in the wildtype-GFP:wildtype-DsRed control experiment did not deviate over time, with approximately equal numbers of each strain observable in the swarm over the course of eight hours. The Δ*ids*-GFP:wildtype-DsRed co-swarm did not show measurable changes until well after swarm emergence (Figure 5), indicating that territorial exclusion did not occur in the swarm inoculum. However, large decreases in the GFP/DsRed ratio were observed in the subsequent consolidation periods between rounds of swarming, starting in the first consolidation phase. Later swarms often contained no observable Δ*ids*-GFP cells.

Reasoning that Ids-mediated exclusion was correlated with consolidation in *P. mirabilis* swarms, we generated “hyperswarming” wild-type and Δ*ids* strains (named “wildtype_-Hs_” and “Δ*ids*_-Hs_,” respectively) that continually swarm outwards without consolidation (31); this leads to rapid surface coverage. We performed swarm-based territorial exclusion assays to test whether hyperswarming protected the Δ*ids*-derived strains from exclusion by wild-type. We observed that neither Δ*ids*_-Hs_:wildtype_-Hs_ nor Δ*ids*_-Hs_:wildtype co-swarms resulted in the exclusion of the hyperswarming Δ*ids* strain (supplementary Figure 5). However, Δ*ids*:wildtype_-Hs_ co-swarms, in which the Δ*ids* strain enters consolidation, did result in territorial exclusion of Δ*ids*-derived cells (Supplementary Figure 5). Outside of the consolidation phase, the Δ*ids*-derived cells that received non-self signals were not effectively excluded, indicating that Ids mismatch does not affect swarm performance in hyperswarming cells. Rather, territorial exclusion was increasingly effective over the course of the co-swarm with initial equal ratios of cells (Figure 5), resulting in the Δ*ids* cells being excluded from the leading edges of swarming colonies.

## Conclusions

Here we expand on models of kin discrimination (5, 8) by showing that the Ids system encompasses a complex and subtle recognition that is attuned to the challenges of rapid migration as a collective along a hard surface. Ids-mediated recognition controls the spatial location of non-self cells over the lifetime of a swarm. It appears that access to a social behavior is impeded via a non-lethal mechanism: the Ids self-recognition selectively induces non-self cells into a lifestyle incompatible with cooperative swarming. Intriguingly, Ids-like proteins are encoded within the genomes of other members of the *Morganellaceae* family, including the pathogen *Providencia stuartii*, suggesting that this mechanism might be more broadly found. Further, these data suggest a model for Ids territorial exclusion in mixed swarms (Figure 6). IdsD is primarily transferred during active swarming when the secretion machinery is produced (21, 25, 39). During consolidation phase, the presence of IdsD in the absence of a cognate IdsE (resulting in unbound IdsD in recipient cells) causes a shift into a distinct transcriptional state (Figure 1) that is partially due to activation of the stringent response via elevated ppGpp levels (Figure 3). This shifted state also causes a phenotype in affected cells that allows increased antibiotic tolerance (Figure 2). We propose that Ids mismatch functions by diverting cells from re-entry into *P. mirabilis’* swarm-consolidation cycle, which results in individual non-self cells being iteratively winnowed out of the migrating swarm front when initially present in equal ratios (Figure 6).

Several potential models could explain exactly how unbound IdsD affects the recipient cell. Our preferred model is that the presence of unbound IdsD in a recipient cell interrupts an essential checkpoint in the differentiation from a swarmer to consolidated cell. While the transcriptomic data provides a reasonable starting point, the list of differences for each Ids mismatch condition as compared to wildtype is quite large. While many genes had a decreased levels of transcripts, several also had increased transcript levels, including multiple fimbriae. These changes do not resemble those previously described during swarm-consolidation transitions (25), suggesting that Ids mismatch induces entry into a novel expression state. It is also formally possible that IdsD might accumulate in the membrane over time, leading to a general stress response. However, several pieces of data contradict such a model. First, hyperswarming Δ*ids* cells receiving a non-self signal are motile and able to swarm with wildtype (Supplementary Figure 5), and excluded Δ*ids* cells are still able to grow and divide *in situ* (17). We therefore deem it unlikely that IdsD functions to directly disrupt membrane integrity or metabolic activity, since these would result in loss of motility or reduced growth under our observed conditions. Further, non-self cells are able to escape Ids-mediated territorial exclusion under laboratory conditions: either overexpression of the master flagellar regulator *flhDC*, which abolishes consolidation to form hyperswarmer cells (31), or deletion of either *relA* or *spoT*, which alters ppGpp levels (Figure 3 and Supplementary Figure 5). Recognition signals need flagellar regulation and internal ppGpp levels to be effective, but how these pathways intersect remains to be uncovered. Interpretations are further complicated, because little is published about the stringent response and/or ppGpp activity in *P. mirabilis*.

**Fig. 6.**
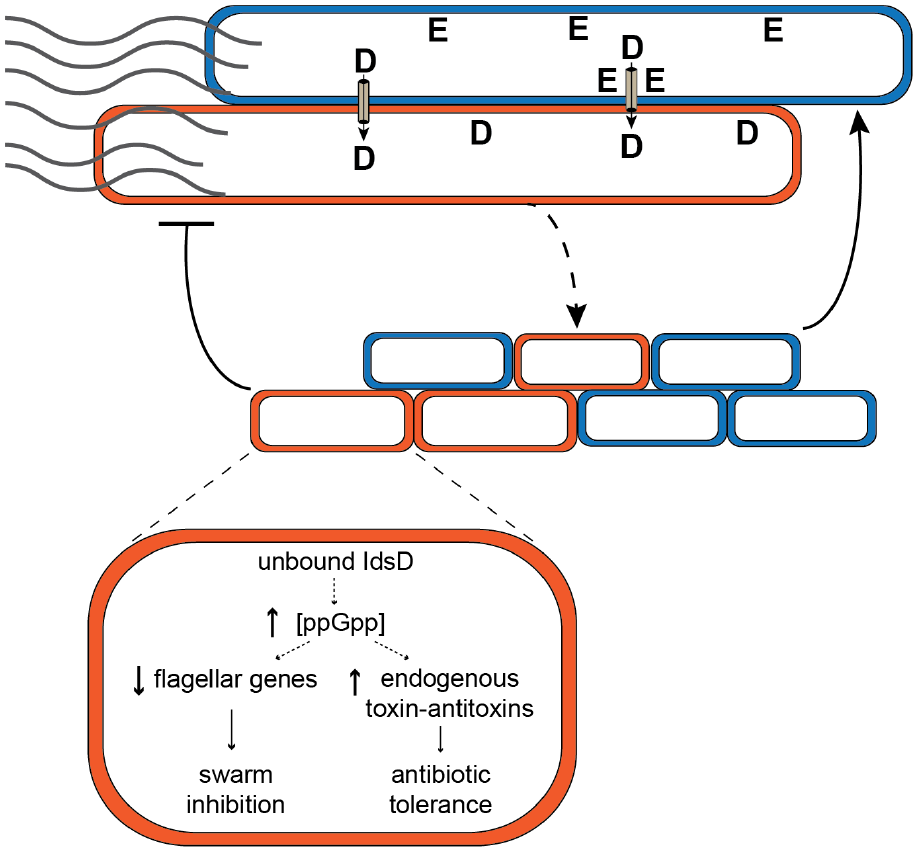
A model for Ids-mediated territorial exclusion. Top, IdsD is exchanged between motile, hyperelongated swarmer cells. Middle, cells expressing a cognate IdsE do not experience any effects from incoming IdsD (green) and progress normally through consolidation phase, forming new swarms. Cells without a cognate IdsE (red) continue to swarm normally after IdsD transfer until entry into consolidation phase. Bottom, upon differentiation into consolidation phase, unbound IdsD induces elevated ppGpp levels and profound transcriptional changes that results in a distinct transcriptional state, rather than differentiation into new swarmer cells. The resultant phenotype exhibits secondary effects including increased antibiotic tolerance.

There are several transcriptional changes in the distinct swarm-incompatible state that are not readily explained by the ppGpp response. We anticipate that ongoing studies into the emergence of bacterial dormancy and related phenotypes (46–49) in other species will help to untangle the order and hierarchy of the Ids-induced changes described here. Several candidate pathways for further analysis are apparent from the transcriptomics datasets. The role of the signaling molecule c-di-GMP in regulating motile/sessile lifestyle changes in many bacteria is well-studied and an attractive target for future work (50). The SOS response, mediated by RecA, has been implicated in persister formation in *E. coli* K12 (51) and may also play a role here. Ids gene regulation, in general, has been linked with the MrpJ transcriptional network important for *P. mirabilis* virulence (30). The observed increased in transcripts for MR/P fimbria (Figure 1)suggests a potential link between Ids-induced changes, MrpJ, and changes in virulence.

More generally, we have presented evidence of a peer-pressure system for recognition that iteratively winnows nonself cells from participating in the collective behavior of swarming—a social activity between cells that is observed among many bacterial species. Ids-mediated macroscale territorial behavior emerges from the sum of cell-cell contacts in a swarm (14, 17, 20, 52). In this Ids model, cells do not receive any information about population composition and behavior other than that from their immediate neighbors, which is different when compared to other examples of bacterial collective behavior. For example, in bacterial quorum sensing, secretion of diffusible small molecules into the environment provides a global tracker accessible to all individuals in a group (53, 54). Therefore, each individual cell has potentially equal access to the external signal, because the signal molecule can freely diffuse between cells. Each *P. mirabilis* cell in a swarm, however, has access only to the signal of a physically adjacent cell. As such, any information about the swarming population as a whole is decentralized and distributed among every member of the swarm in Ids mismatch-mediated exclusion. Access to that information is restricted to clusters of adjacent, neighboring cells. In these respects, Ids-mediated control represents an orthogonal model for collective behavior in bacteria that provides new opportunities to explore cell-cell communication, especially as regards to spatial coordination. Any theoretical model of Ids-mediated behavior will likely need to differ from those describing quorum sensing of diffusible molecules.

The Ids self-recognition system has distinct qualities from other contact-associated systems, which have been described as bacterial communication. CDI systems (55), for example, have been described as lethal (56) or inducing permanent persister-like states (57). However, broad analysis of spatial and temporal dynamics has yet to be pursued, perhaps because CDI generally acts in bulk liquid cultures. *P. mirabilis* swarms could provide an excellent experimentally tractable framework for directly analyzing individual cells before, during, and after Ids communication and for examining the global spatial consequences to these local interactions. Moreover, the ability of the Ids system to temporally and spatially control non-self cells by altering cell state raises the question of which other mechanisms for contact-mediated signaling are present in bacteria to enable sophisticated interactions between individuals.

## Methods

### Media

All strains used in this study are described in Table 1. *P. mirabilis* strains were maintained on LSW-agar (58). CM55 blood agar base agar (Oxoid, Basingstoke UK) was used as a swarm-permissive agar. *E. coli* strains were maintained on Lennox lysogeny broth (LB) agar. All liquid cultures were grown in LB broth at 37°C with shaking. Swarm plates were grown either at room temperature or at 37°C. Antibiotics were added when appropriate at the following concentrations: kanamycin 35 μg ml^−1^, chloramphenicol 50 μg ml^−1^, carbenicillin 100 μg ml^−1^, ampicillin 100 μg ml^−1^, tetracycline 15 μg ml^−1^.

### Strain construction

All chromosomal mutants in BB2000 and the Δ*ids* strain were made as described in (17) with the following modifications for strains constructed *de novo* in this study: the suicide vector was (62) and the conjugative *E. coli* strain was strain MFDpir (61). The *BB2000_0531* transcriptional reporter strain MJT02 includes a gene encoding the Venus fluorescent protein (44) immediately following the stop codon of gene *BB2000_0531*. All chromosomal mutations were confirmed by PCR amplification followed by Sanger sequencing of the amplified product (Genewiz, South Plainfield NJ).

### Co-swarming experiments

Strains were grown overnight at 37°C in LB broth with appropriate antibiotics. Overnight cultures were diluted in LB broth to an optical density at 600 nm (OD_600_) of 1.0, then mixed to the desired experimental ratio and inoculated with an inoculation needle onto a CM55 swarm agar plate. Plates were incubated at 37°C for 18 hours, ensuring that the swarm had covered most of the agar plate. After incubation, swarm composition was measured by using a 48-pin multiblot replicator to sample the swarm and replica plate on non-swarming LSW-agar with relevant antibiotics as described in (14).

### RNA-Seq

Strains were grown on swarm-permissive agar plates with appropriate antibiotics at 37°C. For consolidating cell samples of wildtype, swarm colonies were left to progress overnight and confirmed to be in consolidation phase by light microscopy. Cells from the swarm edge were then harvested by scraping with a plastic loop into 1 ml of RNA Protect solution (Qiagen, Hilden, Germany). The Δ*idsE* and CCS02 samples were harvested after overnight incubation by scraping whole colonies into 1 ml RNA Protect solution. Total RNA was isolated using a RNeasy Mini kit (Qiagen, Hilden, Germany) according to the manufacturer’s instructions. RNA purity was measured using an Agilent 2200 Tapestation (Agilent, Santa Clara, CA). To enrich mRNA, rRNA was digested using terminator 5’ phosphate dependent exonuclease (Illumina, San Diego, CA) according to the manufacturer’s instructions. Enriched RNA samples were purified by phenol-chloroform extraction (63).

The cDNA libraries were prepared from mRNA-enriched RNA samples using an NEBNext Ultra RNA library prep kit (New England Biolabs, Ipswich, MA) according to the manufacturer’s instructions. Libraries were sequenced on an Illumina HiSeq 2500 instrument with 250 bp single-end reads, and base-calling was done with Illumina CASAVA 1.8 in the Harvard University Bauer Core Facility. Sequences were matched to BB2000 reference genome PMID: 24009111 (accession number CP004022) using TopHat2 using default arguments (64). Differential expression data were generated using the Cufflinks RNA-Seq analysis suite (65) run on the Harvard Odyssey cluster. Specifically, the mRNA abundance data were generatedusing Cufflinks 2.1.1 with max-multiread-fraction 0.9 and - multi-read-correct. Samples were combined using cuffmerge with default arguments. Differential expression data were generated using Cuffdiff 2.1.1 with total-hits-norm. The data was analyzed using the CummeRbund package for R and Microsoft Excel. Gene functions were taken from the KEGG and COG databases (66, 67). The data shown in this paper represent the combined analysis of two independent biological repeats and will be available from NCBI GEO.

**Table 1.**
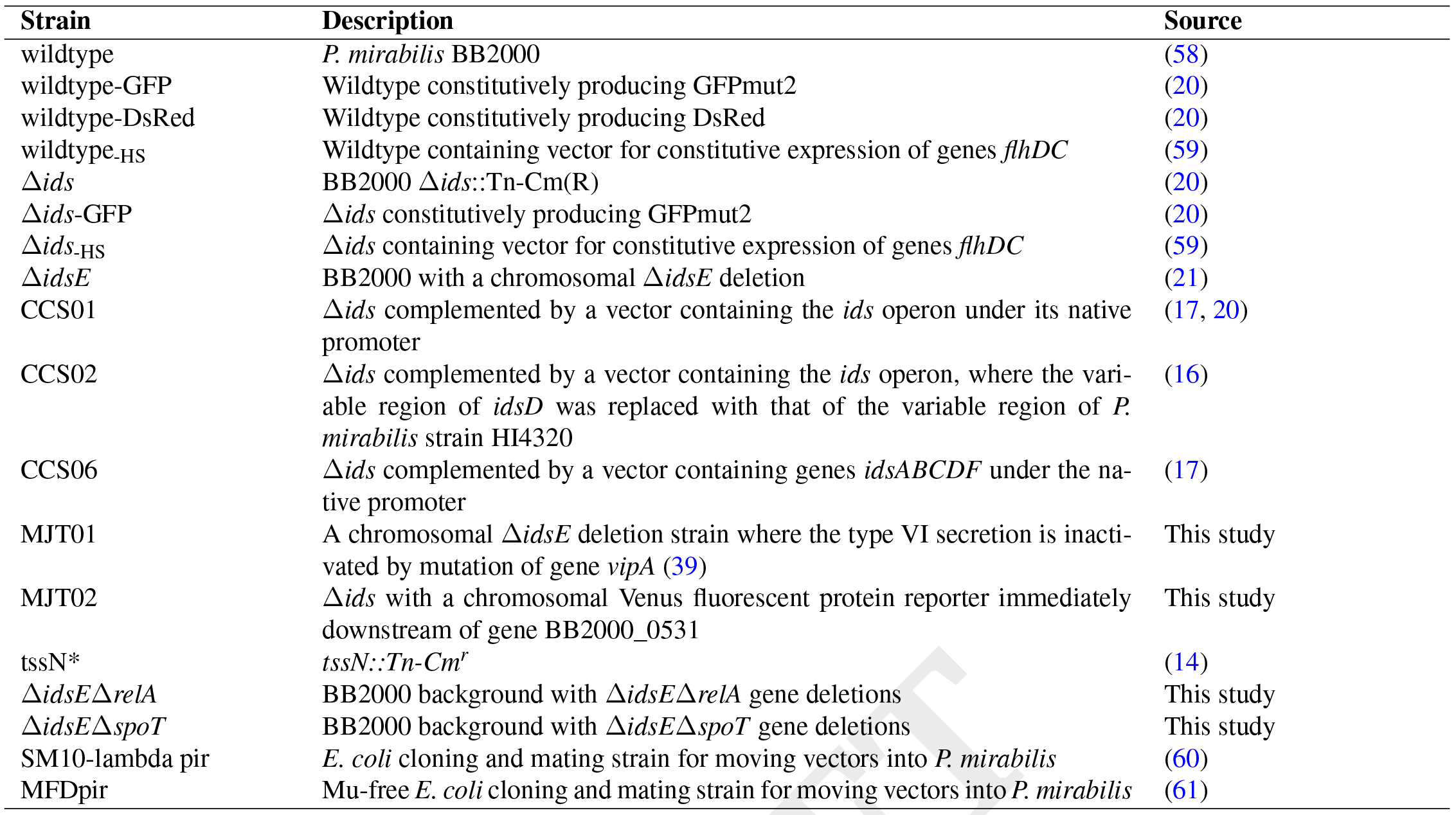
Strains used in this study. All strains are derived from *P. mirabilis* BB2000 unless otherwise noted.

### Fluorescent-activated cell sorting

Samples of excluded Δ*ids* cells were obtained through fluorescent-activated cell sorting (FACS). Fluorescent strains of wildtype-DsRed and Δ*ids*-GFP were grown in liquid at 37°C overnight and normalized to OD_600_ 1.0. Cultures were then mixed to the desired experimental ratio and spotted on swarm agar. After the emergence of the third swarm ring, swarm colonies were harvested into PBS and sorted using a BD FACSAria cell sorter (BD Biosciences, San Jose, CA) into RNA-Protect solution. cDNA samples for RNA-Seq were prepared from sorted samples as described above.

### Microscopy

For experiments where strain ratio during swarming was examined, liquid cultures were grown overnight with shaking at 37°C. Cultures were normalized to OD_600_ 1.0, mixed to the desired experimental ratio, then used to inoculate swarm-permissive agar plates, which were incubated at room temperature overnight at room temperature to allow inoculum development. Plates were then imaged at 30-minute intervals, incubating at 37°C between measurements. For experiments with the *BB2000_0531* transcriptional reporter, 1 μl of mixed, normalized overnight culture were used to inoculate a 1-mm swarm agar pad, which was incubated at 37°C for four hours prior to imaging. Images were taken in GFP (150 ms exposure), RFP (500 ms exposure), and phase contrast channels using a Leica DM5500B microscope (Leica Microsystems, Buffalo Grove IL) and CoolSnap HQ CCD camera (Photometrics, Tucson AZ) cooled to -20°C. MetaMorph version 7.8.0.0 (Molecular Devices, Sunnyvale CA) was used for image acquisition, and FIJI (68) was used for image analysis.

### Antibiotic tolerance assay

Strains were grown on swarm-permissive agar plates at 37°C until swarms reached the second round of consolidation (approximately six hours). Swarm colonies were harvested into LB broth and diluted in LB broth to OD_600_ 1.0. Prior to antibiotic exposure, a sample was taken, serially diluted in LB broth and plated on LSW-agar to count colony-forming units (CFUs/ml) in the sample. Antibiotics were added to the normalized culture at the following concentrations: ampicillin 100 μg ml^−1^, kanamycin 60 μg ml^−1^, streptomycin 50 μg ml^−1^, and ciprofloxacin 1 μg ml^−1^. Each mixture was incubated with shaking at 37°C. At the specified time-points, samples were taken, serially diluted in LB broth, and plated on LSW-agar to measure CFUs/ml. LSW-agar plates were incubated for 16 hours or until visible colonies appeared. Colonies were counted using FIJI. Experiments were performed in triplicate.

### Growth recovery assay following territorial exclusion

Co-swarms of Δ*ids*-GFP and wildtype-DsRed were inoculated onto CM55 plates and allowed to swarm at 37°C. Samples were harvested from swarm plates and sorted via FACS as described above, except that cells were sorted into PBS solution. Immediately after sorting, portions of sorted cell suspension for each strain, containing equal numbers of sorted particles, were used as inoculum for overnight cultures grown at 37°C with shaking in a Tecan Infinite 200 Pro microplate reader (Tecan, Männedorf, Switzerland). OD_600_ measurements were taken hourly. Experiments were performed in triplicate. For experiments with strains CCS01 and CCS02, cell sorting was unnecessary. Instead, clonal swarms were directly harvested into PBS. The resulting suspension was diluted to OD_600_ 0.1 and used as inoculum for cultures containing relevant antibiotics.

### ppGpp quantification using high pressure liquid chromatography

A high-performance liquid chromatography (HPLC)-based method was used to quantify ppGpp levels, based on the work of Varik et al (42). Swarm colonies of *P. mirabilis* were grown to the second swarm ring on CM55 agar. Samples for chromatography were obtained by harvesting cells in 1 ml 1 M acetic acid and immediately flash-freezing in liquid nitrogen. Samples were thawed on ice for 1 hour 30 minutes with occasional vortexing, freeze dried overnight and resuspended in 200 μl MQ-H2O, and then centrifuged at 4°C for 30 min to remove any insoluble fragments. Supernatants were run on a Spherisorb strong ion exchange chromatography column (80 Å, 4.6 by 150 mm, 5 μm, Waters, Milford MA). An isocratic program was used with flow rate 1.5 ml/min in running buffer consisting of 0.36 M ammonium dihydrogen phosphate, 2.5% acetonitrile (v/v), pH 3.6. Nucleotide concentrations were quantified by measuring UV absorbance at 252 nm, comparing peaks to those obtained from purified nucleotide and ppGpp samples (Trilink Biotechnologies, San Diego, CA).

## Supporting information

## ACKNOWLEDGEMENTS

Ian Maynor performed exploratory assays that inspired the experiments described in Figure 5. Edna Stewart performed the experiment described in Supplementary Figure 5. We thank members of the Gibbs, Losick, Gaudet and D'Souza groups for invaluable discussions, the Harvard University Bauer Core Facility for assistance with RNA-Seq and FAS Research Computing for assistance with RNA-Seq analysis. We thank Steven Biller and Polina Kehayova for comments on earlier drafts. This research was funded by the David and Lucile Packard Foundation, the George W. Merck fund, and Harvard University. We declare no conflicts of interest. We have used the HenriquesLab bioRxiv template, developed by Ricardo Henriques, for drafting the preprint version archived at http://www.bioRxiv.org.

## AUTHOR CONTRIBUTIONS

M.J.T. and K.A.G. conceived and coordinated the study, and wrote the paper. M.J.T. performed all experiments except for the experiment described in Supplementary Figure 5 and analyzed all data.

