## Supplementary material for "Peer pressure from a *Proteus mirabilis* self-recognition system controls participation in cooperative swarm motility"

DRAFT

Keywords:

self recognition, *Proteus mirabilis*, kin selection, swarm motility, stringent response, antibiotic tolerance, bacterial communities, social evolution, cell-cell communication, sociomicrobiology

Running title: Ids controls access to swarming

*NB: All materials and methods are described in the main text.*

*The page count continues from the main text.*

*RNA-Seq tables are found on pages 18 - 46.*

### Supplementary Note 1: Supplemental figures

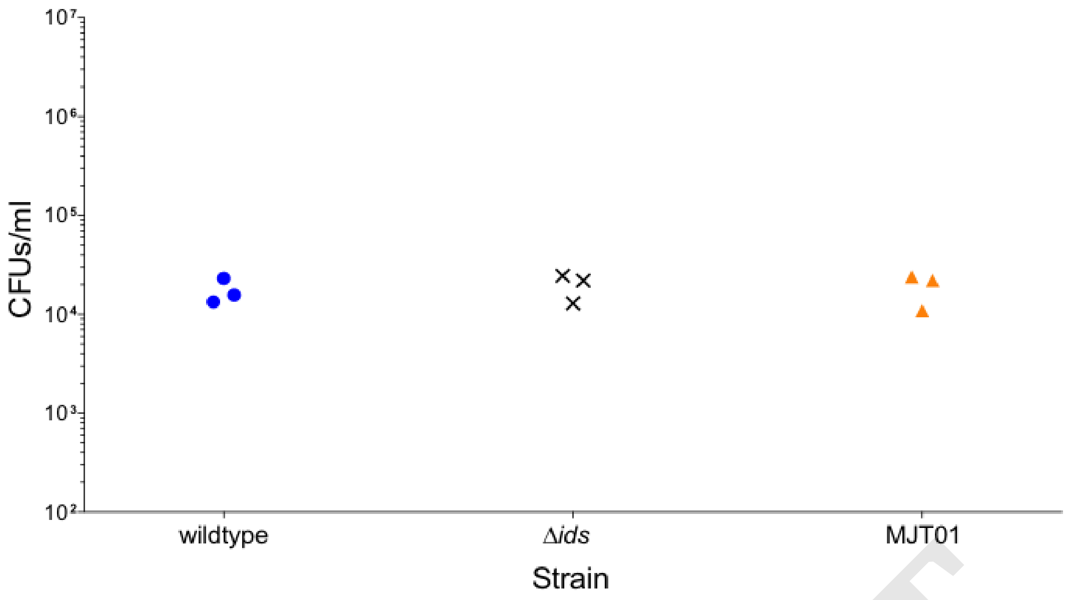


**Fig. SF1. Disruption of the Type VI secretion system removes Ids-mediated antibiotic tolerance.** Cells were harvested from clonal swarm plates and exposed to 100 μg ml$^{-1}$ ampicillin. After 12 hours, viable cells were calculated from growth on fresh LSW- agar. Strains shown are the wildtype, Δ*ids*, and MJT01 strains. Strain MJT01 is an Δ*idsE*-derived strain containing a chromosomal mutation that disrupts its type VI secretion system (39). Each of three biological replicates is presented.

DRAFT

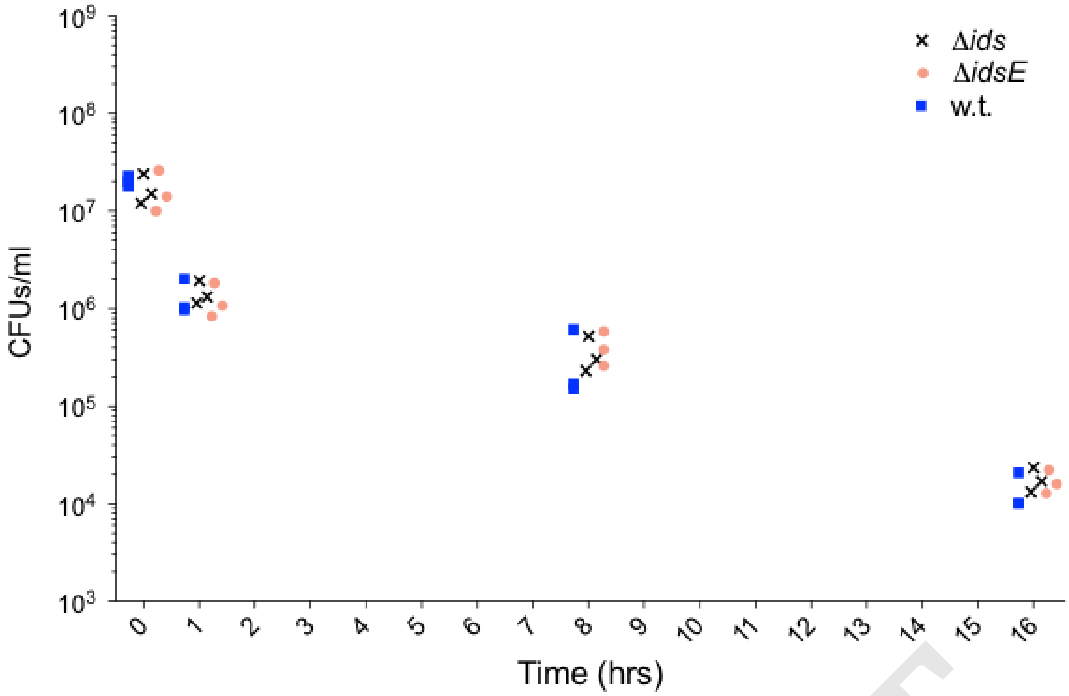


**Fig. SF2. Growth in liquid disrupts Ids-mediated antibiotic tolerance.** Killing curve of the Δ*ids* (black cross), Δ*idsE* (orange triangle), and wildtype (blue square) strains. Cells were isolated from stationary phase cultures and then exposed to 100 μg $ml^{-1}$ ampicillin at each timepoint. Viable cells were calculated after growth on fresh LSW- agar. Each of three biological replicates is presented. Wildtype is labeled "w.t."

DRAFT

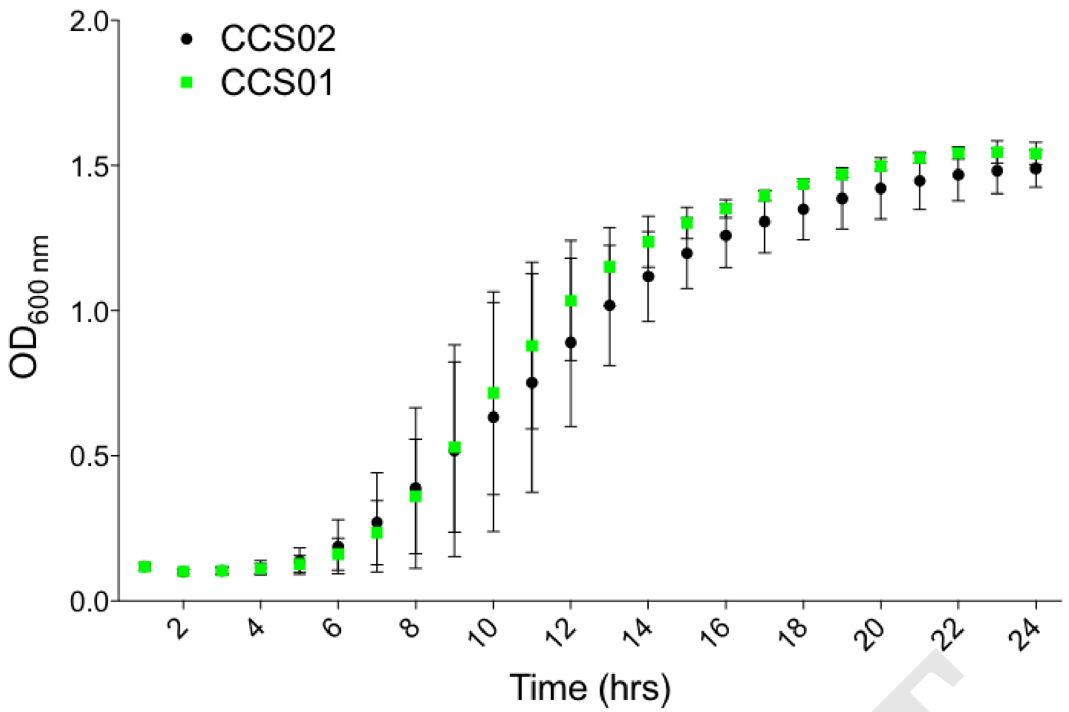


**Fig. SF3. Clonal Ids mismatch mutants do not have long-term growth defects.** Cells were harvested from swarm plates and inoculated into fresh LB media with antibiotics. Growth at 37ºC was assessed by optical density at 600 nm ($OD_{600}$). Strains CCS02, which is an Ids mismatch strain, expresses an IdsE protein unable to bind the incoming IdsD proteins. Strain CCS01 is a Δ*ids* strain complemented by a vector encoding the native *ids* operon and is considered a clonal self population. Three biological repeats were performed. Error bars are standard deviations.

DRAFT

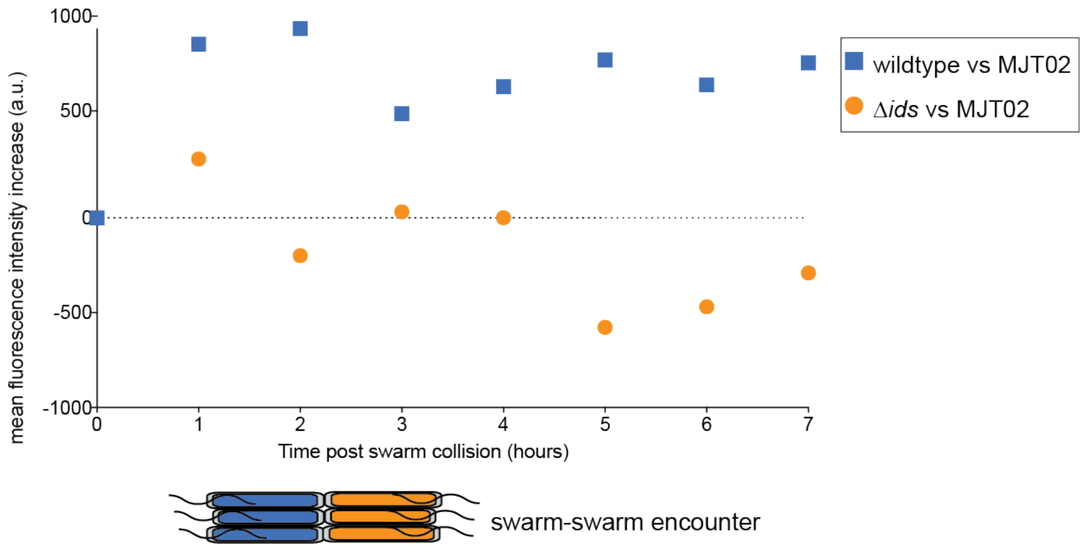


**Fig. SF4. The transcriptional shift observed in cells experiencing Ids mismatch during swarm-swarm encounters occurs prior to visual boundary formation.** A time-course graph showing mean swarm fluorescence intensity over time for two conditions: Δ*ids* carrying a chromosomal BB2000_0531 fluorescence reporter (MJT02, green ovals) swarmed towards wildtype (MJT02:wildtype, blue) or the Δ*ids* strain (MJT02:Δ*ids*, orange). Three biological repeats were performed. Fluorescence was measured over equivalent areas for each experimental condition. Error bars are standard deviations; a.u. means arbitrary units.

DRAFT

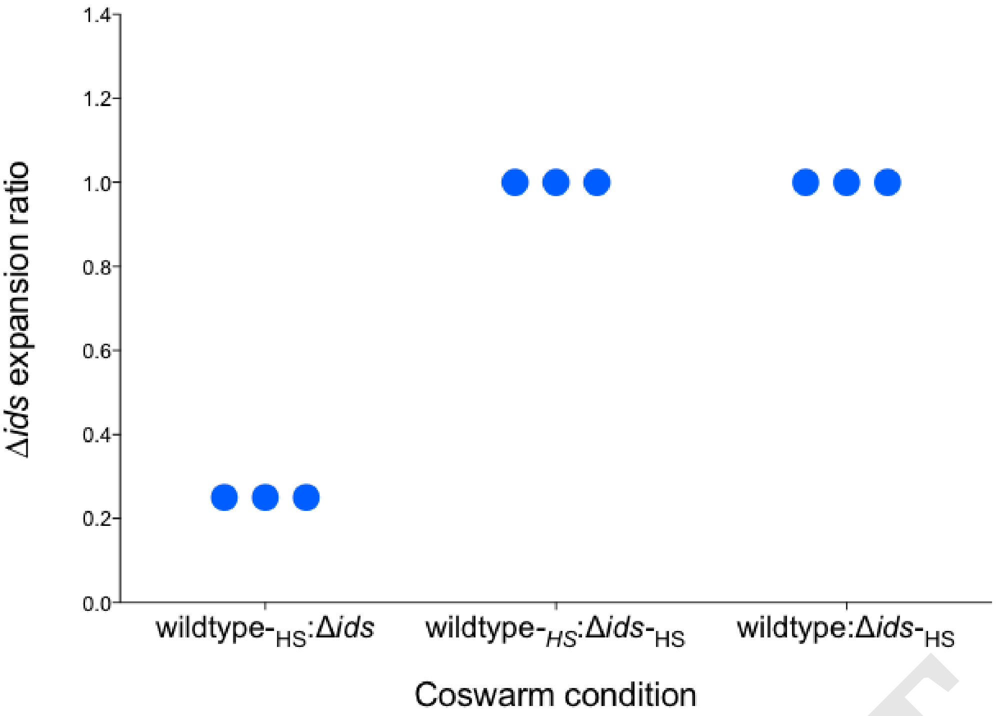


**Fig. SF5. Hyperswarming Δ*ids* escapes Ids-mediated territorial exclusion.** Co-swarms were inoculated at the indicated ratios of wildtype and the Δ*ids* strain as well as derived variants that do not consolidate (i.e., hyperswarm). Formation of hyperswarmer, denoted with a subscript "HS," lineages was due to constitutive in trans expression of *flhDC* as previously described (31). (Left) The Δ*ids* strain was coswarmed with wildtype-HS. (Middle) The Δ*ids*$_{\text{-HS}}$ strain was coswarmed with wildtype$_{\text{-HS}}$. (Right) The Δ*ids*$_{\text{-HS}}$ strain was coswarmed with wildtype$_{\text{-HS}}$. Co-swarms were assessed for spatial distribution as detailed in the main text. Y-axis shows maximum swarm distance of the Δ*ids* strain as a ratio of the swarm distance for the wildtype (right) or wildtype-derived strains (left and middle). Each of three biological replicates is presented.

DRAFT

**Table S1.** Significantly differentially regulated genes between wildtype and clonal Δ*ids*.

| $log_2$—fold change | BB2000 gene name | product |
|---|---|---|
| 8.97778 | BB2000_3003, BB2000_3004, BB2000_3005, BB2000_3006, BB2000_3007 | Ids proteins |
| 2.37131 | BB2000_3145 | 50S ribosomal protein |
| 2.31615 | BB2000_2655 | short hypothetical protein |
| 2.25321 | BB2000_0110 | short hypothetical protein |
| 1.5191 | BB2000_2815 | hypothetical protein |
| -2.68561 | BB2000_1880 | glucose dehydrogenase |

DRAFT

**Table S2.** Significantly differentially regulated genes between CCS01 and CCS06.

| $log_2$-fold change | BB2000 gene name | product |
|---|---|---|
| -7.62439 | BB2000_0139 | hypothetical protein |
| -7.52894 | BB2000_0110 | hypothetical protein |
| -7.00039 | idrA | IdrA |
| -6.79654 | zapD | type I secretion outer membrane protein |
| -6.79654 | zapC | type I secretion protein |
| -6.79654 | zapB | Type I secretion ATP-binding protein |
| -6.79654 | zapA | metalloprotease |
| -6.79654 | zapE | metalloprotease |
| -6.79654 | BB2000_0432 | metalloprotease |
| -6.79654 | BB2000_0433 | metalloprotease |
| -6.70601 | BB2000_1555 | hypothetical protein |
| -6.4165 | ndpA | nucleoid-associated protein NdpA |
| -6.4165 | BB2000_0942 | hypothetical protein |
| -6.20813 | gshA | glutamate–cysteine ligase |
| -6.14722 | nagC | N-acetylglucosamine regulatory protein |
| -6.09512 | mgtC | Mg(2+) transport ATPase protein C |
| -6.09512 | BB2000_1771 | hypothetical protein |
| -6.09512 | BB2000_1772 | hypothetical protein |
| -6.07724 | BB2000_0138 | tetrapyrrole methylase |
| -5.95337 | BB2000_1050 | hypothetical protein |
| -5.95337 | BB2000_1594 | hypothetical protein |
| -5.95337 | fliE | flagellar hook-basal body complex protein |
| -5.91701 | BB2000_0943 | helicase |
| -5.85666 | BB2000_1130 | putative sulfate transporter YchM |
| -5.85666 | fumC | fumarate hydratase |
| -5.81357 | BB2000_1316 | hypothetical protein |
| -5.78566 | holD | DNA polymerase III subunit psi |
| -5.78566 | rplM | 50S ribosomal protein L13 |
| -5.7783 | fliA | flagellar biosynthesis sigma factor |
| -5.65649 | BB2000_0232 | hypothetical protein |
| -5.39259 | BB2000_0858 | CoA-binding protein |
| -5.23493 | flgN | flagella synthesis protein |
| -5.16936 | BB2000_1925 | hypothetical protein |
| -5.14772 | gltX | glutamyl-tRNA synthetase |
| -5.13845 | BB2000_1967 | alpha-2-macroglobulin-like lipoprotein (endopeptidase inhibitor) |
| -5.13845 | intB | prophage integrase |
| -5.13845 | rpsB | 30S ribosomal protein S2 |
| -5.13845 | BB2000_0595 | esterase |
| -5.12694 | adhC | alcohol dehydrogenase (glutathione-dependent formaldehyde dehydrogenase) |
| -5.06481 | sucA | 2-oxoglutarate dehydrogenase E1 component |
| -5.06481 | sucB | dihydrolipoamide succinyltransferase component of 2-oxoglutarate dehydrogenase complex |
| -5.06481 | sucC | succinyl-CoA synthetase subunit beta |
| -5.06481 | sucD | succinyl-CoA synthetase alpha chain |
| -5.06481 | atpI | F0F1 ATP synthase subunit I |
| -5.06481 | pheS | phenylalanyl-tRNA synthetase alpha chain |
| -5.06481 | ihfA | integration host factor subunit alpha |
| -5.05064 | alaS | alanyl-tRNA synthetase |
| -5.0021 | ogt | methylated-DNA–protein-cysteine methyltransferase |
| -5.0021 | cspE | cold shock protein CspE |
| -5.0021 | BB2000_0974 | hypothetical protein |
| -4.91336 | fabH | 3-oxoacyl-[acyl-carrier-protein] synthase III |
| -4.91336 | fabD | malonyl CoA-acyl carrier protein transacylase |

DRAFT

| | | |
|---|---|---|
| -4.89773 | fabG | 3-ketoacyl-(acyl-carrier-protein) reductase |
| -4.89773 | flhC | transcriptional activator FlhC |
| -4.89773 | BB2000_0972 | lipoprotein |
| -4.89773 | BB2000_0973 | Maf-like protein |
| -4.89773 | BB2000_2655 | hypothetical protein |
| -4.89773 | fliZ | flagella biosynthesis protein FliZ |
| -4.89383 | rpsA | 30S ribosomal protein S1 |
| -4.85221 | prsA | ribose-phosphate pyrophosphokinase |
| -4.85221 | dps | DNA starvation/stationary phase protection protein Dps |
| -4.81885 | ompA | outer membrane protein A |
| -4.79169 | ahpC | alkyl hydroperoxide reductase |
| -4.78788 | atpB | ATP synthase A chain |
| -4.78437 | lrp | leucine-responsive transcriptional regulator |
| -4.78025 | rpsI | 30S ribosomal protein S9 |
| -4.77071 | fliC1 | flagellin 1 |
| -4.76033 | ribH | 6,7-dimethyl-8-ribityllumazine synthase |
| -4.69192 | BB2000_2426 | hypothetical protein |
| -4.65181 | znuA | high-affinity zinc transporter periplasmic component |
| -4.57981 | ci | phage repressor |
| -4.54869 | ptsI | phosphoenolpyruvate-protein phosphotransferase |
| -4.5171 | BB2000_0948 | membrane-associated phosphatase |
| -4.39245 | BB2000_1470 | hypothetical protein |
| -4.38516 | thrS | threonyl-tRNA synthetase |
| -4.38072 | tsf | elongation factor Ts |
| -4.37973 | BB2000_1689 | branched chain amino acid transport protein |
| -4.36883 | arsR | arsenical resistance operon repressor |
| -4.32908 | umoA | upregulatior of flagellar operon (exported protein) |
| -4.26049 | BB2000_0690 | hypothetical protein |
| -4.24049 | budA | alpha-acetolactate decarboxylase |
| -4.23931 | BB2000_2794 | transposase/plasmid-related protein |
| -4.2341 | rpoC | DNA-directed RNA polymerase subunit beta’ |
| -4.18377 | rplJ | 50S ribosomal protein L10 |
| -4.17258 | rplA | 50S ribosomal protein L1 |
| -4.1715 | rplK | 50S ribosomal protein L11 |
| -4.15863 | secE | preprotein translocase subunit SecE |
| -4.15863 | tufB | elongation factor Tu |
| -4.15863 | fusA | elongation factor G (EF-G) |
| -4.15863 | rpsG | 30S ribosomal protein S7 |
| -4.15863 | BB2000_2806 | intracellular sulfur oxidation protein |
| -4.15863 | BB2000_2807 | intracellular sulfur oxidation protein |
| -4.15863 | BB2000_2808 | intracellular sulfur oxidation protein |
| -4.15863 | BB2000_2809 | hypothetical protein |
| -4.15863 | BB2000_1903 | acetyltransferase |
| -4.15863 | BB2000_1904 | lipoprotein |
| -4.15863 | BB2000_1905 | hypothetical protein |
| -4.15863 | BB2000_1906 | hypothetical protein |
| -4.15863 | BB2000_1907 | hypothetical protein |
| -4.15863 | aroC | chorismate synthase |
| -4.15863 | BB2000_1909 | methylase |
| -4.15863 | ppiB | peptidyl-prolyl cis-trans isomerase B (rotamase B) |
| -4.13795 | gltS | sodium/glutamate symport carrier protein |
| -4.13795 | lysS | lysyl-tRNA synthetase |
| -4.13795 | prfB | peptide chain release factor 2 |
| -4.13795 | recJ | single-stranded-DNA-specific exonuclease |
| -4.13795 | dsbC | thiol:disulfide interchange protein |
| -4.13795 | xerD | tyrosine recombinase |
| -4.13795 | groS | 10 Kda chaperonin |

DRAFT

| | | |
|---|---|---|
| -4.13728 | fxsA | membrane protein FxsA (suppressor of F exclusion of phage T7) |
| -4.12557 | degS | serine endoprotease |
| -4.12505 | chrR | chromate reductase (NADPH-dependent FMN reductase) |
| -4.12505 | flgB | flagellar basal body rod protein FlgB |
| -4.12505 | BB2000_1770 | hypothetical protein |
| -4.07545 | BB2000_1773 | hypothetical protein |
| -4.07104 | pyrC | dihydroorotase |
| -4.06999 | bssS | biofilm formation regulatory protein BssS |
| -4.06997 | rcsA | colanic acid capsular biosynthesis activation protein (LuxR-family transcriptional regulator) |
| -4.06997 | aroQ | 3-dehydroquinate dehydratase |
| -4.04508 | cfa | cyclopropane fatty acyl phospholipid synthase |
| -4.00702 | fldA | flavodoxin 1 |
| -4.00702 | flgM | anti-sigma28 factor FlgM |
| -3.98086 | asnC | asparaginyl-tRNA synthetase |
| -3.96389 | BB2000_0157 | putative ABC transporter ATP-binding protein |
| -3.94153 | BB2000_0725 | probable transporter |
| -3.93312 | minE | cell division topological specificity factor MinE |
| -3.93312 | minD | cell division inhibitor MinD |
| -3.90589 | atpH | F0F1 ATP synthase subunit delta |
| -3.904 | BB2000_0710 | MFS-family transporter |
| -3.904 | BB2000_2819 | methyl-accepting chemotaxis protein |
| -3.85925 | rluB | 23S rRNA pseudouridylate synthase B |
| -3.85706 | nusB | transcription antitermination protein NusB |
| -3.85706 | cstA | carbon starvation protein |
| -3.85706 | BB2000_1097 | hypothetical protein |
| -3.84328 | BB2000_1100 | fimbrial chaperone |
| -3.8431 | BB2000_1101 | fimbrial protein |
| -3.8179 | BB2000_1102 | fimbrial subunit |
| -3.80326 | BB2000_1103 | fimbrial protein |
| -3.80228 | BB2000_1104 | fimbrial protein |
| -3.75719 | mioC | flavodoxin |
| -3.75582 | ydgA | hypothetical protein |
| -3.75582 | BB2000_2388 | oxidoreductase |
| -3.75582 | icd | isocitrate dehydrogenase |
| -3.75582 | accD | acetyl-CoA carboxylase subunit beta |
| -3.75582 | budB | acetolactate synthase |
| -3.75582 | BB2000_2967 | iron ABC transporter, substrate-binding protein |
| -3.75582 | BB2000_1436 | conjugated bile acid hydrolase |
| -3.75582 | lpp | major outer membrane lipoprotein (murein-lipoprotein) |
| -3.73617 | BB2000_1584 | transcriptional regulator |
| -3.72467 | BB2000_0021 | hypothetical protein |
| -3.72252 | adk | adenylate kinase |
| -3.71905 | BB2000_1956 | lipoprotein |
| -3.7133 | rpmH | 50S ribosomal protein L34 |
| -3.71034 | hpcR | homoprotocatechuate degradative operon repressor (MarR family transcriptional regulator) |
| -3.70446 | BB2000_2855 | signal sensing protein |
| -3.70262 | glpF | glycerol uptake facilitator protein |
| -3.69822 | atpA | F0F1 ATP synthase subunit alpha |
| -3.69439 | BB2000_1717 | hypothetical protein |
| -3.69257 | BB2000_0214 | hypothetical protein |
| -3.67375 | gmk | guanylate kinase |
| -3.67304 | cspB | cold shock protein |
| -3.67304 | BB2000_1718 | hypothetical protein |
| -3.66895 | BB2000_2866 | lipoprotein |

DRAFT

| | | |
|---|---|---|
| -3.66758 | BB2000_1466 | hypothetical protein |
| -3.65291 | atpG | F0F1 ATP synthase subunit gamma |
| -3.58975 | apt | adenine phosphoribosyltransferase |
| -3.57451 | sdaA | L-serine deaminase 1 (L-serine deaminase 1) |
| -3.57285 | BB2000_1540 | aldose 1-epimerase |
| -3.54524 | ribA | GTP cyclohydrolase II |
| -3.5395 | BB2000_0879 | hypothetical protein |
| -3.48619 | emrR | transcriptional repressor MprA |
| -3.48454 | rimI | ribosomal-protein-alanine N-acetyltransferase |
| -3.45608 | BB2000_2516 | LysR-family transcriptional regulator |
| -3.43835 | BB2000_1918 | fimbrial adapter |
| -3.42453 | BB2000_1920 | hypothetical protein |
| -3.4238 | phsA | thiosulfate reductase precursor |
| -3.42174 | phsC | thiosulfate reductase cytochrome b subunit |
| -3.3915 | accB | biotin carboxyl carrier protein |
| -3.3915 | ftsB | cell division protein FtsB |
| -3.36562 | sixA | phosphohistidine phosphatase |
| -3.34679 | pyrF | orotidine-5'-phosphate decarboxylase |
| -3.34439 | BB2000_1334 | tetratricopeptide repeat protein |
| -3.34439 | BB2000_1335 | hypothetical protein |
| -3.33523 | bcr | bicyclomycin/multidrug efflux system |
| -3.33523 | BB2000_0946 | hypothetical protein |
| -3.33523 | gpt | xanthine-guanine phosphoribosyltransferase |
| -3.33523 | gltA | type II citrate synthase |
| -3.33523 | flgC | flagellar basal-body rod protein |
| -3.31976 | tesB | acyl-CoA thioesterase |
| -3.30618 | rnpA | ribonuclease P |
| -3.28734 | frr | ribosome recycling factor |
| -3.28467 | pabA | para-aminobenzoate synthase component II |
| -3.27167 | BB2000_0047 | hypothetical protein |
| -3.27028 | cbpA | curved DNA-binding protein CbpA |
| -3.27028 | aroK | shikimate kinase I |
| -3.27028 | gst | glutathionine S-transferase |
| -3.27028 | udk | uridine kinase |
| -3.27028 | BB2000_0057 | peptidase |
| -3.26396 | ddg | cold-induced palmitoleoyl transferase |
| -3.26396 | BB2000_1964 | hypothetical protein |
| -3.26393 | sfcA | malate dehydrogenase |
| -3.25276 | BB2000_0775 | hypothetical protein |
| -3.25114 | BB2000_1029 | hypothetical protein |
| -3.24303 | dnaA | chromosomal replication initiator protein |
| -3.23123 | BB2000_2738 | serine protease |
| -3.23018 | BB2000_0243 | hypothetical protein |
| -3.20469 | pyrH | uridylate kinase |
| -3.18766 | trmD | tRNA (guanine-N1)-methyltransferase |
| -3.17968 | folA | dihydrofolate reductase |
| -3.16648 | BB2000_1041 | hypothetical protein |
| -3.1656 | uspF | universal stress protein F |
| -3.12757 | fabF | 3-oxoacyl-(acyl carrier protein) synthase II |
| -3.12443 | BB2000_1211 | anaerobic dimethyl sulfoxide reductase chain C |
| -3.1229 | BB2000_0307 | hypothetical protein |
| -3.10776 | apaG | ApaG |
| -3.0856 | fabA | 3-hydroxydecanoyl-(acyl carrier protein) dehydratase |
| -3.08101 | BB2000_0850 | hypothetical protein |
| -3.06119 | fkpA | FKBP-type peptidyl-prolyl cis-trans isomerase |
| -3.05892 | cyoA | cytochrome o ubiquinol oxidase subunit II |
| -3.0251 | BB2000_2076 | hypothetical protein |

DRAFT

| | | |
|---|---|---|
| -3.01739 | rpsT | 30S ribosomal protein S20 |
| -3.00841 | fliL | flagellar basal body-associated protein FliL |
| -2.98301 | hemB | delta-aminolevulinic acid dehydratase |
| -2.97777 | rluD | 23S rRNA pseudouridine synthase D |
| -2.97132 | BB2000_0534 | hypothetical protein |
| -2.95374 | clpB | protein disaggregation chaperone |
| -2.94654 | ispD | 2-C-methyl-D-erythritol 4-phosphate cytidylyltransferase |
| -2.93703 | BB2000_0041 | transposase |
| -2.92666 | BB2000_0171 | RTX-family protein |
| -2.86974 | hupB | DNA-binding protein HU-beta |
| -2.86974 | ppiD | peptidyl-prolyl cis-trans isomerase (rotamase D) |
| -2.86974 | BB2000_0282 | competence protein |
| -2.86974 | BB2000_0283 | hypothetical protein |
| -2.86974 | BB2000_0284 | hypothetical protein |
| -2.86974 | rimM | 16S rRNA-processing protein RimM |
| -2.86974 | uspE | universal stress protein UspE |
| -2.86575 | fnr | fumarate/nitrate reduction transcriptional regulator |
| -2.86514 | BB2000_1698 | hypothetical protein |
| -2.84702 | exbD | biopolymer transport protein |
| -2.84702 | exbB | biopolymer transport protein |
| -2.84702 | gloA | lactoylglutathione lyase |
| -2.84019 | BB2000_2194 | phosphosugar-binding regulatory protein |
| -2.82923 | metK | S-adenosylmethionine synthetase |
| -2.81552 | BB2000_1293 | hypothetical protein |
| -2.8151 | lpdA | dihydrolipoamide dehydrogenase |
| -2.78386 | BB2000_1204 | hypothetical protein |
| -2.7747 | accC | biotin carboxylase |
| -2.7747 | BB2000_1016 | cold shock protein |
| -2.77314 | fliF | flagellar MS-ring protein |
| -2.75984 | fliI | flagellum-specific ATP synthase |
| -2.75713 | fliJ | flagellar biosynthesis chaperone |
| -2.74589 | fliK | flagellar hook-length control protein |
| -2.73892 | BB2000_1549 | lipoprotein |
| -2.73624 | emrE | methyl viologen resistance protein (ethidium resistance protein) |
| -2.73216 | ipk | 4-diphosphocytidyl-2-C-methyl-D-erythritol kinase |
| -2.73216 | lolB | outer membrane lipoprotein LolB |
| -2.73216 | dusB | tRNA-dihydrouridine synthase B |
| -2.73216 | BB2000_2493 | hypothetical protein |
| -2.73216 | BB2000_0591 | putative metalloprotease |
| -2.73216 | uspG1 | universal stress protein G |
| -2.72347 | BB2000_1478 | hypothetical protein |
| -2.72035 | BB2000_2138 | hypothetical protein |
| -2.72035 | uraA | uracil transporter |
| -2.72035 | upp | uracil phosphoribosyltransferase |
| -2.71567 | flgG | flagellar basal-body rod protein (distal rod protein) |
| -2.7133 | BB2000_0905 | hypothetical protein |
| -2.71226 | modB | molybdate ABC transporter permease protein |
| -2.70419 | hns | DNA-binding protein (histone-like structuring protein) |
| -2.70419 | BB2000_1915 | hypothetical protein |
| -2.68083 | BB2000_0532 | outer membrane protein assembly complex subunit YfiO |
| -2.68083 | rpoE | RNA polymerase sigma factor RpoE |
| -2.68083 | BB2000_1537 | hypothetical protein |
| -2.68083 | msrB | peptide methionine sulfoxide reductase |
| -2.68083 | fadL | long-chain fatty acid outer membrane transporter |
| -2.68083 | thiD | phosphomethylpyrimidine kinase |
| -2.66727 | BB2000_2820 | methyl-accepting chemotaxis protein |
| -2.66727 | BB2000_2944 | LacI-family transcriptional regulator |

DRAFT

| | | |
|---|---|---|
| -2.664 | BB2000_2945 | hypothetical protein |
| -2.664 | BB2000_2946 | hypothetical protein |
| -2.65407 | BB2000_2948 | hypothetical protein |
| -2.65183 | BB2000_2949 | dihydrodipicolinate synthase-family protein |
| -2.65156 | BB2000_2950 | hypothetical protein |
| -2.65156 | BB2000_0792 | hypothetical protein |
| -2.64773 | rplS | 50S ribosomal protein L19 |
| -2.64171 | hpr | phosphohistidinoprotein-hexose phosphotransferase component of PTS system (Hpr) |
| -2.64171 | uspA | universal stress protein A |
| -2.63155 | BB2000_0146 | serine/threonine transporter SstT |
| -2.61527 | efp | elongation factor P |
| -2.61469 | lpxD | UDP-3-O-[3-hydroxymyristoyl] glucosamine N-acyltransferase |
| -2.61193 | ptsN | PTS IIA-like nitrogen-regulatory protein PtsN |
| -2.60876 | sthA | soluble pyridine nucleotide transhydrogenase |
| -2.58066 | flgF | flagellar basal-body rod protein |
| -2.5767 | BB2000_1968 | hypothetical protein |
| -2.56561 | BB2000_1963 | lipoprotein |
| -2.56536 | acrR | DNA-binding transcriptional repressor AcrR |
| -2.56536 | BB2000_0297 | cytoplasmic sulphur reductase |
| -2.56536 | kefA | potassium efflux protein KefA |
| -2.56536 | BB2000_0938 | hypothetical protein |
| -2.56536 | cysS | cysteinyl-tRNA synthetase |
| -2.56536 | BB2000_2598 | radical SAM superfamily protein |
| -2.56536 | narP | nitrate/nitrite response regulator |
| -2.56415 | idrB | IdrB |
| -2.56245 | BB2000_0308 | putative GTP-binding protein YjiA |
| -2.55841 | BB2000_0873 | hypothetical protein |
| -2.54851 | BB2000_1383 | hypothetical protein |
| -2.53444 | pyrD | dihydroorotate dehydrogenase |
| -2.53386 | BB2000_1449 | hypothetical protein |
| -2.5311 | BB2000_1679 | lipid kinase |
| -2.51191 | aldB | aldehyde dehydrogenase |
| -2.47086 | BB2000_1792 | hypothetical protein |
| -2.46225 | BB2000_1793 | acetyltransferase |
| -2.45942 | BB2000_1794 | NADH-dependent flavin oxidoreductase |
| -2.45874 | BB2000_1795 | hypothetical protein |
| -2.45723 | BB2000_2844 | transposase |
| -2.45723 | BB2000_2050 | hypothetical protein |
| -2.45255 | tpx | thiol peroxidase |
| -2.44042 | epd | erythrose 4-phosphate dehydrogenase |
| -2.43555 | glpT | sn-glycerol-3-phosphate transporter |
| -2.43555 | BB2000_2321 | hypothetical protein |
| -2.43555 | nlpB | lipoprotein |
| -2.43126 | BB2000_0757 | hypothetical protein |
| -2.42977 | macA | macrolide transporter subunit MacA |
| -2.42959 | macB | macrolide transporter ATP-binding /permease protein |
| -2.42486 | clpS | ATP-dependent Clp protease adaptor protein |
| -2.42304 | clpA | ATP-dependent Clp protease ATP-binding subunit |
| -2.41867 | rraA | ribonuclease activity regulator protein RraA |
| -2.41395 | speG | spermidine N(1)-acetyltransferase (diamine acetyltransferase) |
| -2.41068 | dnaN | DNA polymerase III subunit beta |
| -2.40031 | BB2000_0476 | metallo-beta-lactamase superfamily protein |
| -2.39614 | BB2000_1576 | glucose 1-dehydrogenase |
| -2.38836 | BB2000_2175 | hypothetical protein |
| -2.38719 | BB2000_2176 | hypothetical protein |

DRAFT

| | | |
|---|---|---|
| -2.38717 | BB2000_2177 | ArsR-family transcriptional regulator |
| -2.37226 | spr | putative outer membrane lipoprotein |
| -2.37122 | BB2000_0951 | elongation factor P-like protein |
| -2.3648 | BB2000_0953 | MutT/NUDIX family protein |
| -2.36211 | nfo | endonuclease IV |
| -2.36058 | BB2000_0955 | transposase |
| -2.35577 | BB2000_0950 | hypothetical protein |
| -2.35577 | glpQ | glycerophosphodiester phosphodiesterase |
| -2.34351 | cyoE | protoheme IX farnesyltransferase |
| -2.34073 | BB2000_0611 | hypothetical protein |
| -2.33788 | tktA | transketolase |
| -2.33449 | BB2000_1459 | AsnC-family transcriptional regulator |
| -2.33283 | BB2000_0038 | hypothetical protein |
| -2.32491 | pabB | para-aminobenzoate synthase component I |
| -2.32048 | BB2000_1696 | hypothetical protein |
| -2.31604 | dgkA | diacylglycerol kinase |
| -2.31399 | BB2000_2548 | pyridoxal-dependent decarboxylase |
| -2.29621 | BB2000_2549 | Mg(2+)/citrate complex transporter |
| -2.27728 | deoB | phosphopentomutase |
| -2.27158 | deoA | thymidine phosphorylase |
| -2.26571 | deoC | deoxyribose-phosphate aldolase |
| -2.26571 | BB2000_2554 | Na+ dependent nucleoside transporter |
| -2.26571 | BB2000_2555 | TatD-related deoxyribonuclease |
| -2.24943 | BB2000_1360 | dsRNA-binding protein |
| -2.24943 | BB2000_1361 | hypothetical protein |
| -2.24943 | BB2000_1362 | hypothetical protein |
| -2.24943 | mipA | MltA-interacting protein precursor |
| -2.24943 | hypF | hydrogenase maturation protein |
| -2.24943 | dadB | alanine racemase, catabolic |
| -2.24943 | dadA | D-amino acid dehydrogenase small subunit |
| -2.2345 | BB2000_2885 | hypothetical protein |
| -2.23393 | BB2000_2886 | plasmid-related protein |
| -2.20559 | aroB | 3-dehydroquinate synthase |
| -2.20559 | ptsO | phosphocarrier protein |
| -2.20458 | BB2000_0088 | hypothetical protein |
| -2.19252 | pykA | pyruvate kinase |
| -2.18899 | hexR | DNA-binding transcriptional regulator HexR |
| -2.18837 | tolB | translocation protein TolB |
| -2.18676 | pal | peptidoglycan-associated outer membrane lipoprotein |
| -2.17162 | BB2000_0653 | hypothetical protein |
| -2.17162 | rplT | 50S ribosomal protein L20 |
| -2.17162 | flgK | flagellar hook-associated protein 1 |
| -2.16662 | lexA | LexA repressor |
| -2.15842 | btuC | vtamin B12-transporter permease |
| -2.15842 | btuD | vitamin B12 import ATP-binding protein |
| -2.15842 | arnB | UDP-4-amino-4-deoxy-L-arabinose–oxoglutarate aminotransferase |
| -2.15842 | arnC | undecaprenyl phosphate 4-deoxy-4-formamido-L-arabinose transferase |
| -2.15842 | arnA | bifunctional UDP-glucuronic acid decarboxylase/UDP-4-amino-4-deoxy-L-arabinose formyltransferase |
| -2.15842 | BB2000_1083 | polysaccharide deacetylase |
| -2.15842 | arnT | 4-amino-4-deoxy-L-arabinose transferase |
| -2.15842 | BB2000_1085 | hypothetical protein |
| -2.15842 | BB2000_1086 | hypothetical protein |
| -2.15842 | BB2000_0398 | hypothetical protein |
| -2.15842 | fliD | flagellar capping protein |

DRAFT

| | | |
|---|---|---|
| -2.14383 | fliS | flagellar protein FliS |
| -2.14383 | fliT | flagella protein |
| -2.14383 | fabZ | (3R)-hydroxymyristoyl-ACP dehydratase |
| -2.14383 | glnS | glutaminyl-tRNA synthetase |
| -2.14383 | mdtK | multidrug efflux protein |
| -2.14022 | BB2000_2310 | oligo-nucleotide binding protein (suppressor of ushA transcription) |
| -2.13881 | tgt | queuine tRNA-ribosyltransferase |
| -2.1244 | cheZ | chemotaxis regulator CheZ |
| -2.12014 | BB2000_2623 | hypothetical protein |
| -2.11964 | cheY | chemotaxis response regulator |
| -2.11356 | BB2000_0376 | hypothetical protein |
| -2.10765 | cueR | MerR-family transcriptional regulator (copper efflux regulator) |
| -2.09846 | pgk | phosphoglycerate kinase |
| -2.09758 | BB2000_0108 | cytochrome d ubiquinol oxidase subunit III |
| -2.09758 | BB2000_1027 | hypothetical protein |
| -2.09758 | BB2000_1460 | LysE-type transporter |
| -2.09003 | BB2000_3262 | hypothetical protein |
| -2.05317 | BB2000_0706 | acetyltransferase |
| -2.04992 | BB2000_2260 | hypothetical protein |
| -2.03484 | caiD | carnitinyl-CoA dehydratase |
| -2.02839 | BB2000_0286 | AsnC-family transcriptional regulator |
| -2.01944 | BB2000_1524 | hypothetical protein |
| -2.01214 | gntX | gluconate metabolism protein |
| -2.00073 | glpE | thiosulfate sulfurtransferase (gluconate metabolism protein) |
| -1.99776 | glpG | intramembrane serine protease GlpG |
| -1.99762 | glpR | DNA-binding transcriptional repressor GlpR |
| -1.99241 | metF | 5,10-methylenetetrahydrofolate reductase |
| -1.99127 | corC | magnesium and cobalt efflux protein |
| -1.98928 | BB2000_2854 | insulinase (Peptidase family M16) |
| -1.96765 | rbsC | ribose ABC transporter permease protein |
| -1.96678 | rbsA | D-ribose transporter ATP binding protein |
| -1.96627 | rbsD | high affinity ribose transport protein |
| -1.94616 | BB2000_0747 | hypothetical protein |
| -1.94616 | terE | tellurite resistance protein |
| -1.92467 | terB | tellurite resistance protein |
| -1.91238 | terZ | tellurite resistance protein |
| -1.91238 | idrD | IdrD |
| -1.90014 | focA | probable formate transporter |
| -1.89467 | BB2000_0779 | hypothetical protein |
| -1.88659 | ppx | exopolyphosphatase |
| -1.8865 | accA | acetyl-CoA carboxylase carboxyltransferase subunit alpha |
| -1.88624 | BB2000_2273 | hypothetical protein |
| -1.8819 | BB2000_1829 | phage antitermination protein |
| -1.87908 | glyQ | glycyl-tRNA synthetase subunit alpha |
| -1.87817 | BB2000_0738 | glutathione S-transferase |
| -1.86296 | dcd | deoxycytidine triphosphate deaminase |
| -1.85934 | cutC | copper homeostasis protein CutC |
| -1.85443 | BB2000_1135 | chaperone |
| -1.84853 | ompF | outer membrane porin |
| -1.8455 | maeB | malic enzyme |
| -1.84089 | BB2000_0317 | TetR-family transcriptional regulator |
| -1.83993 | ssb | single-strand binding protein |
| -1.83813 | kefB | glutathione-regulated potassium-efflux system protein KefB |
| -1.83813 | kefG | glutathione-regulated potassium-efflux system ancillary protein KefG |
| -1.83813 | recF | recombination protein F |

DRAFT

| | | |
|---|---|---|
| -1.83813 | mreB | rod shape-determining protein MreB |
| -1.83813 | mreC | rod shape-determining protein MreC |
| -1.83813 | mreD | rod shape-determining protein MreD |
| -1.82119 | BB2000_0077 | inhibitor of septum formation |
| -1.81957 | cafA | ribonuclease G |
| -1.81957 | BB2000_0079 | hypothetical protein |
| -1.81957 | BB2000_0080 | carbon-nitrogen hydrolase |
| -1.81957 | tldD | protease TldD |
| -1.81957 | BB2000_0082 | exported ribonuclease |
| -1.81653 | BB2000_0083 | ribonuclease inhibitor |
| -1.80129 | thiB | thiamine ABC transporter, substrate-binding protein |
| -1.80099 | thiP | thiamine transporter membrane protein |
| -1.79931 | thiQ | thiamine transporter ATP-binding subunit |
| -1.7925 | BB2000_2464 | Possible excisionase |
| -1.79092 | BB2000_2465 | hypothetical protein |
| -1.78825 | rapA | ATP-dependent helicase HepA |
| -1.78621 | rpoS | RNA polymerase sigma factor RpoS |
| -1.78468 | can | carbonic anhydrase |
| -1.78468 | BB2000_0326 | hypothetical protein |
| -1.77257 | BB2000_1694 | hypothetical protein |
| -1.77257 | wzz | ferric enterobactin transport protein FepE |
| -1.77257 | BB2000_1844 | lipoprotein |
| -1.77079 | BB2000_1292 | beta-eliminating lyase |
| -1.76717 | dcuC | C4-dicarboxylate transporter DcuC |
| -1.76642 | BB2000_2098 | Z-ring-associated protein |
| -1.75415 | gcvR | glycine cleavage system transcriptional repressor |
| -1.75003 | bcp | thioredoxin-dependent thiol peroxidase |
| -1.74268 | purN | phosphoribosylglycinamide formyltransferase (5'-phosphoribosylglycinamide transformylase) |
| -1.74235 | BB2000_2088 | hypothetical protein |
| -1.74235 | tpiA | triosephosphate isomerase |
| -1.73782 | dksA | DnaK transcriptional regulator DksA |
| -1.73782 | BB2000_2229 | fimbrial subunit |
| -1.73782 | BB2000_1892 | hypothetical protein |
| -1.73782 | folC | bifunctional folylpolyglutamate synthase/ dihydrofolate synthase |
| -1.73141 | secF | preprotein translocase subunit SecF |
| -1.72278 | BB2000_2389 | hypothetical protein |
| -1.72278 | pepB | aminopeptidase B |
| -1.72278 | BB2000_0145 | hypothetical protein |
| -1.72088 | dut | deoxyuridine 5'-triphosphate nucleotidohydrolase |
| -1.72003 | BB2000_0869 | LysR-family transcriptional regulator |
| -1.70778 | srmB | ATP-dependent RNA helicase SrmB |
| -1.70778 | BB2000_0048 | hypothetical protein |
| -1.70458 | potD | spermidine/putrescine ABC transporter periplasmic substrate-binding protein |
| -1.70232 | potC | spermidine/putrescine ABC transporter membrane protein |
| -1.70116 | potB | spermidine/putrescine ABC transporter membrane protein |
| -1.69845 | potA | putrescine/spermidine ABC transporter ATPase protein |
| -1.69527 | dapA | dihydrodipicolinate synthase |
| -1.69395 | purU | formyltetrahydrofolate deformylase |
| -1.69106 | rseA | anti-RNA polymerase sigma factor SigE |
| -1.68596 | BB2000_1438 | peptidoglycan-binding protein |
| -1.68552 | sufE | cysteine desulfuration protein |
| -1.68552 | sufS | bifunctional cysteine desulfurase/selenocysteine lyase |
| -1.68552 | sufD | cysteine desulfurase activator complex subunit SufD |
| -1.68495 | sufC | cysteine desulfurase ATPase component |

DRAFT

| | | |
|---|---|---|
| -1.68254 | sufB | cysteine desulfurase activator complex subunit SufB |
| -1.65913 | sufA | scaffold protein for iron-sulfur cluster assembly |
| -1.64243 | BB2000_1446 | thioesterase |
| -1.63433 | pbpC | penicillin-binding protein 1C |
| -1.63433 | cmk | cytidylate kinase |
| -1.63347 | rnt | ribonuclease T |
| 1.50771 | dppF | dipeptide transporter ATP-binding subunit |
| 1.50771 | dppD | dipeptide transporter ATP-binding subunit |
| 1.50874 | BB2000_1229 | integrase/recombinase |
| 1.51063 | BB2000_2275 | hypothetical protein |
| 1.51063 | BB2000_2276 | hypothetical protein |
| 1.51063 | BB2000_2277 | hypothetical protein |
| 1.51063 | BB2000_2278 | hypothetical protein |
| 1.51276 | BB2000_2972 | multidrug efflux protein (MFS-family transporter) |
| 1.51999 | BB2000_2336 | isochorismatase |
| 1.52472 | clcA | chloride channel protein |
| 1.52642 | BB2000_2478 | hypothetical protein |
| 1.52819 | BB2000_1703 | universal stress protein |
| 1.53151 | fpr | ferredoxin-NADP reductase |
| 1.53473 | BB2000_2216 | arylsulfatase |
| 1.53514 | idsE3 | IdsE3 |
| 1.53514 | idsF2 | IdsF2 |
| 1.53624 | BB2000_0923 | phage protein |
| 1.54262 | bioB | biotin synthase |
| 1.54262 | bioF | 8-amino-7-oxononanoate synthase |
| 1.54262 | bioC | biotin synthesis protein BioC |
| 1.54262 | bioD | dithiobiotin synthetase |
| 1.55214 | hcr | HCP oxidoreductase, NADH-dependent |
| 1.55382 | BB2000_3133 | hypothetical protein |
| 1.55731 | BB2000_2761 | hypothetical protein |
| 1.56085 | BB2000_2169 | hypothetical protein |
| 1.56509 | BB2000_0934 | surface polysaccharide modification acyltransferase |
| 1.57315 | narI | respiratory nitrate reductase 1 gamma chain |
| 1.57581 | glnK | nitrogen regulatory protein P-II |
| 1.57581 | amtB | ammonium transporter |
| 1.58107 | BB2000_0620 | hypothetical protein |
| 1.58107 | BB2000_0621 | hypothetical protein |
| 1.58158 | potE | putrescine transporter |
| 1.58319 | BB2000_0962 | autotransporter |
| 1.58341 | hmuR2 | hemin receptor |
| 1.58353 | BB2000_1820 | phage protein |
| 1.58487 | BB2000_3112 | cellulose synthase regulator protein |
| 1.58662 | BB2000_2747 | hypothetical protein |
| 1.58783 | mltA | murein transglycosylase A |
| 1.59056 | BB2000_1624 | hypothetical protein |
| 6.01507 | BB2000_0381 | ABC transporter, ATP-binding subunit |
| 1.59758 | BB2000_3115 | signaling protein |
| 1.60327 | rluA | ribosomal large subunit pseudouridine synthase |
| 1.60394 | BB2000_2956 | ABC-transporter, permease protein |
| 1.60726 | BB2000_2333 | hypothetical protein |
| 1.60969 | BB2000_1592 | surface polysaccharide modification acyltransferase |
| 1.61377 | BB2000_0624 | hypothetical protein |
| 1.61377 | BB2000_0625 | hypothetical protein |
| 1.61377 | BB2000_0626 | hypothetical protein |
| 1.61392 | aslB | Radical SAM superfamily protein (probable arylsulfatase-activating protein) |
| 1.61605 | cysH | phosphoadenosine phosphosulfate reductase |

DRAFT

| | | |
|---|---|---|
| 1.61605 | cysI | sulfite reductase subunit beta |
| 1.61605 | cysJ | sulfite reductase [NADPH] flavoprotein alpha-component |
| 1.62872 | BB2000_3100 | fimbrial chaperone |
| 1.62925 | lysA | diaminopimelate decarboxylase |
| 1.63542 | BB2000_3096 | multidrug efflux protein |
| 1.63542 | BB2000_3097 | multidrug efflux protein (MFS-family transporter) |
| 1.6372 | BB2000_0424 | hypothetical protein |
| 1.64605 | argG | argininosuccinate synthase |
| 1.64721 | BB2000_2201 | autotransporter |
| 1.64761 | BB2000_0531 | sigma 54 modulation protein |
| 1.65303 | BB2000_0924 | phage protein |
| 1.6541 | BB2000_2696 | type III secretion system protein |
| 1.65852 | fixA | putative electron transfer flavoprotein FixA |
| 1.65922 | BB2000_1226 | efflux protein |
| 1.65939 | poxB | pyruvate dehydrogenase |
| 1.66979 | BB2000_1345 | lipoprotein |
| 1.67495 | hisD | histidinol dehydrogenase |
| 1.67684 | galU | UTP–glucose-1-phosphate uridylyltransferase |
| 1.67764 | BB2000_2660 | hypothetical protein |
| 1.68282 | BB2000_1037 | lipase |
| 1.68584 | BB2000_2228 | hypothetical protein |
| 1.68627 | fdhF | formate dehydrogenase H |
| 1.68753 | caiC | putative crotonobetaine/carnitine-CoA ligase |
| 1.69369 | BB2000_3134 | TonB-dependent receptor |
| 1.69476 | BB2000_0319 | hypothetical protein |
| 1.70222 | BB2000_0550 | MFS-family transporter |
| 1.70895 | BB2000_0549 | GntR-family transcriptional regulator |
| 1.71499 | cat | chloramphenicol acetyltransferase |
| 1.72376 | BB2000_0912 | hypothetical protein |
| 1.73247 | BB2000_1344 | lipoprotein |
| 1.74529 | atfC | outer membrane usher protein |
| 1.74571 | celY | cellulase |
| 1.74942 | agaZ | tagatose 6-phosphate kinase |
| 1.75094 | chbR | DNA-binding transcriptional regulator ChbR |
| 1.75355 | BB2000_0351 | two-component sensor kinase |
| 1.76752 | BB2000_2242 | phage protein |
| 1.76752 | BB2000_2243 | phage protein |
| 1.77009 | BB2000_2258 | phage protein |
| 1.77009 | BB2000_2259 | hypothetical protein |
| 1.77263 | argO | arginine exporter protein |
| 1.77946 | BB2000_3013 | fimbrial protein |
| 1.78956 | BB2000_0745 | transposase |
| 1.79022 | dppB | dipeptide transporter permease DppB |
| 1.79054 | BB2000_3014 | hypothetical protein |
| 1.79409 | BB2000_2339 | sodium:sulfate symporter |
| 1.79437 | csaA | protein secretion chaperone |
| 1.79875 | BB2000_2913 | hypothetical protein |
| 1.79875 | BB2000_2914 | hypothetical protein |
| 1.79875 | BB2000_2915 | hypothetical protein |
| 1.80386 | BB2000_2924 | haemagglutinin |
| 1.8092 | BB2000_1091 | GntR-family transcriptional regulator |
| 1.81084 | phoA | alkaline phosphatase |
| 1.81669 | dmsB | anaerobic dimethyl sulfoxide reductase chain B |
| 1.81771 | BB2000_1014 | hypothetical protein |
| 1.81898 | BB2000_0477 | LysR-family transcriptional regulator |
| 1.81995 | fhlA | formate hydrogenlyase transcriptional activator |
| 1.82128 | caiB | crotonobetainyl-CoA:carnitine CoA-transferase |

DRAFT

| | | |
|---|---|---|
| 1.82329 | hycI | hydrogenase 3 maturation protease |
| 1.82329 | hyfJ | hydrogenase-4 component J |
| 1.82329 | hyfI | hydrogenase-4 component I |
| 1.82329 | hyfH | hydrogenase 4 subunit H |
| 1.82329 | hyfG | hydrogenase-4 component G |
| 1.82329 | hyfF | hydrogenase 4 subunit F |
| 1.82329 | hyfE | hydrogenase 4 membrane subunit |
| 1.82329 | hyfD | hydrogenase 4 subunit D |
| 1.82329 | hyfC | hydrogenase-4 component C |
| 1.82329 | hyfB | hydrogenase 4 subunit B |
| 1.82329 | hyfA | hydrogenase-4 component A |
| 1.82559 | BB2000_2350 | fimbrial subunit |
| 1.82559 | BB2000_2351 | fimbrial subunit |
| 1.83199 | BB2000_2989 | sodium:solute symporter |
| 1.83297 | BB2000_2345 | fimbrial outer membrane usher protein |
| 1.83318 | BB2000_1145 | ABC transporter, ATP-binding protein |
| 1.85725 | caiT | L-carnitine/gamma-butyrobetaine antiporter |
| 1.8674 | BB2000_2919 | aminomethyltransferase |
| 1.87075 | rnz | ribonuclease Z |
| 1.87167 | BB2000_2264 | phage protein |
| 1.87373 | arsB | arsenical pump membrane protein |
| 1.87448 | BB2000_2244 | tail length tape measure protein |
| 1.87592 | BB2000_2600 | outer membrane usher protein |
| 1.89049 | cysG | siroheme synthase |
| 1.90934 | BB2000_1606 | hypothetical protein |
| 1.91529 | BB2000_0352 | two-component response regulator |
| 1.91746 | agaD | N-acetylgalactosamine-specific PTS system, EIID component |
| 1.91746 | agaW | N-acetylgalactosamine-specific PTS system, EIIC component |
| 1.92216 | dhaK1 | dihydroxyacetone kinase (glycerone kinase), kinase subunit |
| 1.92216 | dhaK2 | dihydroxyacetone kinase, phosphatase subunit |
| 1.9247 | BB2000_2694 | type III secretion system protein |
| 1.92884 | BB2000_0916 | hypothetical protein |
| 1.93118 | idsE2 | IdsE2 |
| 1.94082 | BB2000_2226 | fimbrial subunit |
| 1.94351 | BB2000_0384 | TonB-dependent siderophore receptor |
| 1.94714 | BB2000_2503 | hypothetical protein |
| 1.95175 | BB2000_2951 | probable carbohydrate kinase |
| 1.95512 | BB2000_1569 | hypothetical protein |
| 1.95652 | BB2000_1825 | phage protein |
| 1.95754 | mrpH | fimbrial adhesin |
| 1.96457 | BB2000_2325 | LuxR-family transcriptional regulator |
| 1.96986 | BB2000_0389 | substrate-binding protein |
| 1.97202 | BB2000_1495 | fimbrial adhesin |
| 1.97556 | fixB | electron transfer flavoprotein alpha subunit for carnitine metabolism |
| 1.99331 | BB2000_3086 | ABC transporter, substrate-binding protein |
| 1.99358 | BB2000_0551 | hydrolase |
| 1.99486 | BB2000_0918 | hypothetical protein |
| 1.99486 | BB2000_0919 | hypothetical protein |
| 1.99486 | BB2000_0920 | hypothetical protein |
| 1.99486 | BB2000_0921 | phage protein |
| 1.99529 | BB2000_2960 | outer membrane protein |
| 2.00618 | BB2000_2262 | phage lysozyme |
| 2.00618 | BB2000_2263 | phage protein |
| 2.01126 | BB2000_1605 | hypothetical protein |
| 2.01222 | chbC | N,N'-diacetylchitobiose-specific PTS system transporter subunit IIC |

DRAFT

| | | |
|---|---|---|
| 2.01488 | caiB | crotonobetainyl-CoA:carnitine CoA-transferase |
| 2.0221 | BB2000_2354 | fimbrial outer membrane usher protein |
| 2.02402 | agaS | tagatose-6-phosphate ketose/aldose isomerase |
| 2.02473 | BB2000_3111 | cellulose synthase catalytic subunit [UDP-forming] |
| 2.0306 | BB2000_0385 | decarboxylase |
| 2.03312 | fixC | putative oxidoreductase FixC |
| 2.03516 | sufI | repressor protein for FtsI |
| 2.04216 | BB2000_0383 | siderophore biosynthesis protein |
| 2.04267 | cueO | multicopper oxidase |
| 2.05196 | BB2000_1301 | hypothetical protein |
| 2.05196 | BB2000_1302 | hypothetical protein |
| 2.05222 | BB2000_2640 | MFS-family transporter |
| 2.06057 | dcuB | anaerobic C4-dicarboxylate transporter |
| 2.07144 | copA | copper exporting ATPase |
| 2.07542 | narG | respiratory nitrate reductase 1 alpha chain |
| 2.07542 | narH | respiratory nitrate reductase 1 beta chain |
| 2.07542 | narJ | respiratory nitrate reductase 1 delta chain |
| 2.08088 | BB2000_1225 | acetyltransferase |
| 2.08177 | BB2000_0925 | phage protein |
| 2.08177 | BB2000_0926 | phage protein |
| 2.08177 | BB2000_0927 | phage protein |
| 2.08177 | BB2000_0928 | phage protein |
| 2.08613 | fdrA | membrane protein FdrA |
| 2.09853 | leuC | 3-isopropylmalate dehydratase large subunit |
| 2.1009 | BB2000_2238 | phage host specificity protein |
| 2.11332 | BB2000_1798 | hypothetical protein |
| 2.11332 | BB2000_1799 | branched-chain amino acid transporter |
| 2.12178 | BB2000_3093 | amidohydrolase/metallopeptidase |
| 2.12944 | speF | ornithine decarboxylase |
| 2.13091 | BB2000_2912 | transacylase |
| 2.13422 | BB2000_0911 | hypothetical protein |
| 2.1585 | BB2000_1227 | hypothetical protein |
| 2.16719 | BB2000_1583 | ABC transporter, substrate-binding protein |
| 2.1706 | BB2000_2247 | phage protein |
| 2.17336 | BB2000_2916 | ATP-binding protein |
| 2.17336 | BB2000_2917 | beta-ketoacyl-ACP synthase |
| 2.17336 | BB2000_2918 | beta-ketoacyl-ACP synthase |
| 2.18689 | mrpC | fimbrial outer membrane usher protein |
| 2.18839 | BB2000_1493 | aminotransferase |
| 2.19564 | ydgI | arginine/ornithine antiporter |
| 2.2014 | BB2000_3114 | cellulose biosynthesis protein |
| 2.20152 | aceB | malate synthase A |
| 2.20678 | BB2000_0909 | phage endopeptidase (lysis protein) |
| 2.21343 | proW | glycine betaine transporter membrane protein |
| 2.21343 | proV | glycine betaine/L-proline ABC transporter, ATP-binding protein |
| 2.21621 | BB2000_1475 | hypothetical protein |
| 2.21698 | speF | ornithine decarboxylase |
| 2.21864 | BB2000_1339 | transferase |
| 2.22309 | BB2000_1297 | Na+/H+ antiporter |
| 2.22437 | BB2000_2923 | holo-[acyl-carrier protein] synthase |
| 2.2352 | BB2000_3008 | hypothetical protein |
| 2.24032 | BB2000_2661 | LysR-family transcriptional regulator |
| 2.24379 | BB2000_2634 | hypothetical protein |
| 2.24379 | BB2000_2635 | hypothetical protein |
| 2.24379 | BB2000_2636 | hypothetical protein |
| 2.24587 | arcC | carbamate kinase |

DRAFT

| | | |
|---|---|---|
| 2.24739 | leuA | 2-isopropylmalate synthase |
| 2.2493 | BB2000_3085 | ABC transporter permease protein |
| 2.25003 | BB2000_2428 | hypothetical protein |
| 2.25117 | BB2000_1811 | hypothetical protein |
| 2.25956 | BB2000_2348 | peroxidase |
| 2.26889 | argC | N-acetyl-gamma-glutamyl-phosphate reductase |
| 2.30526 | mrpJ | fimbrial operon regulator |
| 2.30802 | BB2000_2250 | phage protein |
| 2.30802 | BB2000_2251 | phage protein |
| 2.30802 | BB2000_2252 | phage protein |
| 2.30997 | BB2000_2684 | chaperone protein |
| 2.33972 | BB2000_2693 | type III secretion system protein |
| 2.35591 | BB2000_2683 | cell invasion protein |
| 2.36644 | BB2000_1223 | hypothetical protein |
| 2.36967 | BB2000_2695 | type III secretion system protein |
| 2.38411 | BB2000_2114 | hypothetical protein |
| 2.38411 | BB2000_2115 | toxin |
| 2.38411 | BB2000_2116 | toxin |
| 2.38411 | BB2000_2117 | toxin |
| 2.39115 | BB2000_3015 | fimbrial protein |
| 2.39577 | BB2000_0933 | phage protein |
| 2.40318 | BB2000_2200 | hypothetical protein |
| 2.40367 | BB2000_2576 | radical SAM superfamily protein |
| 2.41673 | BB2000_2511 | hypothetical protein |
| 2.43067 | BB2000_2334 | hypothetical protein |
| 2.4338 | BB2000_2995 | PTS system, EIIBC component |
| 2.43941 | BB2000_1239 | hypothetical protein |
| 2.44033 | dmsA | dimethyl sulfoxide reductase chain A |
| 2.45218 | nirD | nitrite reductase small subunit |
| 2.45218 | nirB | nitrite reductase [NAD(P)H] large subunit |
| 2.46217 | BB2000_2697 | type III secretion system regulatory protein |
| 2.47564 | leuO | leucine transcriptional activator |
| 2.47982 | ipdC | indole-3-pyruvate decarboxylase |
| 2.4848 | BB2000_0910 | hypothetical protein |
| 2.52112 | BB2000_2633 | hypothetical protein |
| 2.52749 | uca | major fimbrial subunit |
| 2.5278 | dmsB | anaerobic dimethyl sulfoxide reductase chain B |
| 2.52885 | fixB | electron transfer flavoprotein alpha subunit for carnitine metabolism |
| 2.52965 | BB2000_1625 | hypothetical protein |
| 2.53102 | hisG | ATP phosphoribosyltransferase |
| 2.53879 | BB2000_1498 | fimbrial subunit |
| 2.54757 | BB2000_1133 | hypothetical protein |
| 2.55603 | BB2000_1607 | hypothetical protein |
| 2.55603 | BB2000_1608 | hypothetical protein |
| 2.56654 | BB2000_2921 | fatty acyl chain dehydratase |
| 2.5684 | BB2000_2254 | head maturation protease |
| 2.57025 | BB2000_2239 | phage tail protein |
| 2.57025 | BB2000_2240 | phage protein |
| 2.57958 | BB2000_0386 | pyridoxal-phosphate dependent enzyme |
| 2.57958 | BB2000_0387 | octopine/opine/tauropine dehydrogenase |
| 2.57982 | BB2000_0395 | hypothetical protein |
| 2.58916 | BB2000_0913 | hypothetical protein |
| 2.58916 | BB2000_0914 | hypothetical protein |
| 2.58916 | BB2000_0915 | hypothetical protein |
| 2.60024 | BB2000_0929 | phage protein |
| 2.60024 | BB2000_0930 | phage protein |

DRAFT

| | | |
|---|---|---|
| 2.60024 | BB2000_0931 | phage protein |
| 2.60024 | BB2000_0932 | phage protein |
| 2.60317 | BB2000_2911 | hypothetical protein |
| 2.61436 | fadD | long-chain-fatty-acid–CoA ligase |
| 2.62315 | BB2000_2681 | cell invasion protein |
| 2.62367 | BB2000_2501 | demethylmenaquinone methyltransferase |
| 2.66066 | BB2000_0388 | MFS-family transporter |
| 2.66238 | BB2000_1256 | transport protein |
| 2.68747 | BB2000_2255 | phage portal protein |
| 2.68747 | BB2000_2256 | phage terminase, large subunit |
| 2.70356 | BB2000_2269 | hypothetical protein |
| 2.70356 | BB2000_2270 | hypothetical protein |
| 2.71386 | fbpC | ferric transporter ATP-binding subunit |
| 2.72063 | BB2000_1591 | hypothetical protein |
| 2.72724 | dmsA | dimethyl sulfoxide reductase chain A |
| 2.73072 | BB2000_2279 | hypothetical protein |
| 2.73561 | BB2000_2346 | fimbrial chaperone protein |
| 2.73766 | BB2000_2836 | hypothetical protein |
| 2.74133 | BB2000_2353 | fimbrial chaperone protein |
| 2.76965 | BB2000_1496 | fimbrial outer membrane usher protein |
| 2.77722 | BB2000_3110 | hypothetical protein |
| 2.79363 | hcp | hydroxylamine reductase |
| 2.80738 | BB2000_3016 | fimbrial protein |
| 2.80824 | BB2000_0896 | hypothetical protein |
| 2.81596 | caiA | crotonobetainyl-CoA dehydrogenase |
| 2.83572 | BB2000_2248 | phage protein |
| 2.85597 | BB2000_2922 | 3-oxoacyl-[acyl-carrier protein] reductase |
| 2.88101 | fixA | putative electron transfer flavoprotein FixA |
| 2.88776 | BB2000_2249 | phage protein |
| 2.96062 | cysD | sulfate adenylyltransferase subunit 2 |
| 3.05738 | mrpG | fimbrial subunit |
| 3.05968 | caiA | crotonobetainyl-CoA dehydrogenase |
| 3.06275 | BB2000_0340 | hypothetical protein |
| 3.11084 | BB2000_3103 | fimbrial subunit |
| 3.12918 | BB2000_1294 | LysE-family transporter |
| 3.13612 | BB2000_1570 | carbohydrate kinase/trancriptional regulator |
| 3.13692 | BB2000_2343 | fimbrial subunit |
| 3.14704 | BB2000_2355 | fimbrial subunit |
| 3.17389 | narK | nitrite extrusion protein (MFS-family transporter) |
| 3.1739 | BB2000_1283 | MFS-family transporter |
| 3.1739 | BB2000_1284 | hypothetical protein |
| 3.19382 | pmpA | fimbrial subunit |
| 3.20432 | BB2000_1810 | hypothetical protein |
| 3.22269 | BB2000_2352 | fimbrial protein |
| 3.32801 | BB2000_0831 | hypothetical protein |
| 3.33659 | BB2000_2253 | major capsid protein |
| 3.33968 | BB2000_1499 | fimbrial subunit |
| 3.44759 | BB2000_2344 | fimbrial subunit |
| 3.49784 | BB2000_2502 | hypothetical protein |
| 3.5195 | bioA | adenosylmethionine-8-amino-7-oxononanoate aminotransferase |
| 3.52405 | BB2000_2257 | phage terminase, small subunit |
| 3.62037 | BB2000_0830 | hypothetical protein |
| 3.64197 | BB2000_0922 | phage protein |
| 3.64613 | BB2000_1497 | fimbrial chaperone protein |
| 3.99141 | BB2000_1552 | hypothetical protein |
| 4.14668 | BB2000_3158 | hypothetical protein |

DRAFT

| | | |
|---|---|---|
| 4.14668 | BB2000_3159 | hypothetical protein |
| 4.82774 | BB2000_3198 | serine acetyltransferase |

DRAFT

**Table S3.** Significantly differentially regulated genes between CCS01 and CCS06.

| $log_2$-fold change | BB2000 gene name | product |
|---|---|---|
| -4.5289 | BB2000_0427, BB2000_0428, BB2000_0429, BB2000_0430 | zapABCD |
| -4.04955 | BB2000_1015 | lipase |
| -3.7314 | BB2000_1500 | fimbrial operon regulator |
| -3.6785 | BB2000_1717 | hypothetical protein |
| -3.5754 | BB2000_1016 | cold shock protein |
| -3.49282 | flgC | flagellar basal-body rod protein |
| -3.48269 | BB2000_1826 | phage protein (endopeptidase/lysis protein) |
| -3.47585 | ddg | cold-induced palmitoleoyl transferase |
| -3.41201 | rnk | nucleoside diphosphate kinase regulator |
| -3.2631 | fliZ, fliA, BB2000_1711, fliC2 | flagella biosynthesis protein FliZ, flagellar biosynthesis sigma factor, hypothetical protein, hypothetical protein |
| -3.17636 | BB2000_2819 | methyl-accepting chemotaxis protein |
| -3.09508 | fliJ | flagellar biosynthesis chaperone |
| -3.07686 | bfd | bacterioferritin-associated ferredoxin |
| -3.06231 | cspA | cold shock protein |
| -3.05536 | cspB | cold shock protein |
| -3.05119 | BB2000_2949 | dihydrodipicolinate synthase-family protein |
| -3.01576 | flgA | flagella basal body P-ring formation protein |
| -2.9013 | BB2000_2950 | hypothetical protein |
| -2.88481 | BB2000_0879 | hypothetical protein |
| -2.87996 | fliF, fliG, fliH, fliI | flagellar MS-ring protein, flagellar motor switch protein G, flagellar assembly protein H, flagellum-specific ATP synthase |
| -2.8306 | BB2000_1294 | LysE-family transporter |
| -2.75331 | BB2000_1827 | phage protein |
| -2.73139 | BB2000_1381 | outer membrane protein (attachment invasion locus protein) |
| -2.71296 | BB2000_2565 | hypothetical protein |
| -2.66101 | BB2000_3198 | serine acetyltransferase |
| -2.57831 | BB2000_1824 | phage protein |
| -2.57672 | BB2000_1033, map1 | hypothetical protein, methionine aminopeptidase |
| -2.5747 | BB2000_2344 | fimbrial subunit |
| -2.5683 | BB2000_1215, BB2000_1216 | PadR-family transcriptional regulator, hypothetical protein |
| -2.56305 | BB2000_0924 | phage protein |
| -2.52905 | flgD | basal-body rod modification protein |
| -2.52684 | BB2000_1097 | hypothetical protein |
| -2.492 | fumC, BB2000_1316 | fumarate hydratase, hypothetical protein |
| -2.42215 | BB2000_0832 | Rhs-family protein |
| -2.40861 | BB2000_1586 | hypothetical protein |
| -2.36323 | BB2000_1818, BB2000_1819 | phage protein, phage protein |
| -2.3213 | BB2000_1470 | hypothetical protein |
| -2.31577 | BB2000_0342 | transcriptional regulator |
| -2.2976 | flhC | transcriptional activator FlhC |
| -2.26955 | BB2000_2352 | fimbrial protein |
| -2.25484 | BB2000_1828 | phage holin (lysis protein) |
| -2.23795 | pbpC | penicillin-binding protein 1C |
| -2.22757 | ribB | 3, 4-dihydroxy-2-butanone 4-phosphate synthase |
| -2.20865 | BB2000_2855 | signal sensing protein |
| -2.20668 | BB2000_0704 | threonine and homoserine efflux system |
| -2.20651 | BB2000_1829 | phage antitermination protein |
| -2.19849 | pagP, pagQ | palmitoyl transferase |
| -2.19146 | BB2000_1815, BB2000_1816 | phage protein, phage protein |
| -2.18995 | BB2000_1825 | phage protein |
| -2.17953 | BB2000_1809 | hypothetical protein |
| -2.17555 | sthA | soluble pyridine nucleotide transhydrogenase |

DRAFT

| | | |
|---|---|---|
| -2.16621 | hpmB | hemolysin activator protein (two-partner secretion system ac cessory protein) |
| -2.16269 | BB2000_1584 | transcriptional regulator |
| -2.1596 | BB2000_2750 | hypothetical protein |
| -2.15855 | BB2000_0744 | hypothetical protein |
| -2.13634 | terA | tellurite resistance protein |
| -2.11023 | BB2000_1820 | phage protein |
| -2.10422 | flgH | flagellar basal body L-ring protein |
| -2.08224 | fliK | flagellar hook-length control protein |
| -2.05511 | rplV | 50S ribosomal protein L22 |
| -2.05341 | BB2000_3499 | lipoprotein |
| -2.04662 | BB2000_2477 | transposase |
| -2.04396 | flgB | flagellar basal body rod protein FlgB |
| -2.03386 | BB2000_2249 | phage protein |
| -2.02962 | fliE | flagellar hook-basal body complex protein |
| -2.02236 | BB2000_2246 | phage protein |
| -2.01885 | BB2000_3459 | hypothetical protein |
| -2.01871 | flgG | flagellar basal-body rod protein (distal rod protein) |
| -2.01609 | BB2000_2250, BB2000_2251, BB2000_2252 | phage protein, phage protein, phage protein |
| -2.01388 | BB2000_1466 | hypothetical protein |
| -1.95844 | csaA | protein secretion chaperone |
| -1.92888 | BB2000_2664, BB2000_2665 | hypothetical protein, hypothetical protein |
| -1.91794 | acs | acetyl-CoA synthetase |
| -1.89363 | bioA | adenosylmethionine-8-amino-7-oxononanoate aminotrans ferase |
| -1.89049 | BB2000_2274 | hypothetical protein |
| -1.87246 | BB2000_2310 | oligo-nucleotide binding protein (suppressor of ushA transcrip tion) |
| -1.84937 | BB2000_2321 | hypothetical protein |
| -1.84847 | BB2000_0828 | Rhs-family protein |
| -1.84034 | BB2000_2951 | probable carbohydrate kinase |
| -1.82824 | BB2000_0710 | MFS transporter |
| -1.82255 | purE | phosphoribosylaminoimidazole carboxylase catalytic subunit |
| -1.81668 | umoA | upregulator of flagellar operon (exported protein) |
| -1.81097 | rplC | 50S ribosomal protein L3 |
| -1.79966 | BB2000_2948 | hypothetical protein |
| -1.79606 | BB2000_2388 | oxidoreductase |
| -1.76481 | hpaC | 4-hydroxyphenylacetate 3-monooxygenase, reductase compo nent |
| -1.75641 | BB2000_0157 | putative ABC transporter ATP-binding protein |
| -1.71853 | rplB | 50S ribosomal protein L2 |
| -1.71844 | BB2000_3500 | acetyltransferase |
| -1.70908 | BB2000_1292 | beta-eliminating lyase |
| -1.68397 | ccm | membrane protein (Ccm1 protein) |
| -1.67602 | rpsJ | 30S ribosomal protein S10 |
| -1.65258 | BB2000_0156 | TonB-like protein |
| -1.64118 | rplE | 50S ribosomal protein L5 |
| -1.62894 | BB2000_1465 | iron utilization protein |
| -1.61621 | caiE | carnitine operon protein CaiE |
| -1.60387 | rplX | 50S ribosomal protein L24 |
| -1.59794 | emrR | transcriptional repressor MprA |
| -1.57655 | BB2000_1277 | ABC transporter ATP-binding protein |
| -1.57487 | metF | 5, 10-methylenetetrahydrofolate reductase |
| -1.53226 | BB2000_2947 | hypothetical protein |
| -1.52043 | rplN | 50S ribosomal protein L14 |
| 1.54585 | BB2000_3066 | hypothetical protein |

DRAFT

| | | |
|---|---|---|
| 1.55085 | BB2000_0885 | phage replication protein |
| 1.55646 | BB2000_0619 | hypothetical protein |
| 1.57944 | glpQ | glycerophosphodiester phosphodiesterase |
| 1.60273 | BB2000_1494 | fimbrial subunit |
| 1.6642 | umoC | upregulator of flagellar master operon |
| 1.67492 | relE, BB2000_2873 | hypothetical protein |
| 1.75798 | BB2000_1915 | hypothetical protein |
| 1.76336 | bioD | dethiobiotin synthetase |
| 1.78261 | BB2000_1256 | transport protein |
| 1.85284 | BB2000_0966 | hypothetical protein |
| 1.87002 | cpxP | periplasmic protein |
| 1.90918 | BB2000_1900 | hypothetical protein |
| 1.92804 | BB2000_3158 | hypothetical protein |
| 1.92804 | BB2000_3159 | hypothetical protein |
| 1.93113 | hupA | DNA-binding protein HU-alpha (HU-2) |
| 1.95145 | BB2000_1910 | ferritin-like protein |
| 1.95615 | focA | probable formate transporter |
| 1.96074 | oppA | oligopeptide ABC transporter, oligopeptide-binding protein |
| 1.98106 | BB2000_1347 | lipoprotein |
| 1.98458 | BB2000_3206 | hypothetical protein |
| 2.0472 | hyb0 | hydrogenase 2 small subunit |
| 2.04745 | mscL | large-conductance mechanosensitive channel |
| 2.05419 | BB2000_1225 | acetyltransferase |
| 2.05792 | BB2000_3202 | hypothetical protein |
| 2.06765 | BB2000_1476 | hypothetical protein |
| 2.07174 | BB2000_0283 | hypothetical protein |
| 2.07307 | BB2000_3487 | hypothetical protein |
| 2.11788 | BB2000_3399 | hypothetical protein |
| 2.12087 | pspC | DNA-binding transcriptional activator PspC |
| 2.12899 | uca | major fimbrial subunit |
| 2.12959 | BB2000_1346 | lipoprotein |
| 2.13238 | BB2000_1222 | hypothetical protein |
| 2.14666 | BB2000_3230 | hypothetical protein |
| 2.14776 | ftnA | ferritin |
| 2.16803 | BB2000_0223 | hypothetical protein |
| 2.17051 | BB2000_2079 | hypothetical protein |
| 2.17953 | BB2000_1572 | hypothetical protein |
| 2.18264 | holE2 | DNA polymerase III, theta subunit |
| 2.1916 | BB2000_0090 | probable sigma(54) modulation protein |
| 2.21484 | BB2000_2113 | hypothetical protein |
| 2.22212 | BB2000_0850 | hypothetical protein |
| 2.22446 | rob | right origin-binding protein |
| 2.23097 | BB2000_0492 | CsbD family general stress response protein |
| 2.2349 | BB2000_0620 | plasmid stabilization proteins ParE and antitoxin CC2985 |
| 2.24108 | BB2000_2067, BB2000_2068, BB2000_2070 | hypothetical protein, hypothetical protein, hypothetical protein |
| 2.37747 | BB2000_3243 | hypothetical protein |
| 2.39124 | aphA | acid phosphatase/phosphotransferase |
| 2.39207 | tolC | outer membrane channel protein |
| 2.3984 | BB2000_3244 | hypothetical protein |
| 2.42723 | grpE | heat shock protein |
| 2.4299 | BB2000_1552 | hypothetical protein |
| 2.4513 | BB2000_2355 | fimbrial subunit |
| 2.45713 | BB2000_1884 | PTS system EIIA component |
| 2.45877 | BB2000_1140 | lipoprotein |
| 2.47441 | BB2000_2182 | lipoprotein |
| 2.49323 | fdhF | formate dehydrogenase-H, selenopolypeptide subunit |

DRAFT

| | | |
|---|---|---|
| 2.52301 | BB2000_1553 | hypothetical protein |
| 2.52643 | pspA | phage shock protein PspA |
| 2.52742 | BB2000_0891 | hypothetical protein |
| 2.54178 | intB | prophage integrase |
| 2.58607 | pstS | phosphate ABC transporter periplasmic substrate-binding protein PstS |
| 2.58749 | mrpJ | fimbrial operon regulator |
| 2.60486 | asnA | asparagine synthetase AsnA |
| 2.6327 | BB2000_1164 | hypothetical protein |
| 2.65299 | frdA | fumarate reductase flavoprotein subunit |
| 2.65335 | BB2000_0395 | hypothetical protein |
| 2.685 | BB2000_2846 | hypothetical protein |
| 2.74456 | pmfA | major fimbrial subunit |
| 2.77355 | BB2000_0873 | hypothetical protein |
| 2.82506 | BB2000_0330 | hypothetical protein |
| 2.87824 | BB2000_2098 | Z-ring-associated protein |
| 2.88548 | ompW | outer membrane protein W |
| 2.93693 | BB2000_0670 | hypothetical protein |
| 2.95495 | BB2000_1178 | lipoprotein |
| 3.01194 | mrpA | major mannose-resistant/Proteus-like fimbrial protein |
| 3.01521 | BB2000_1970 | hypothetical protein |
| 3.05046 | BB2000_1383, BB2000_1384 | hypothetical protein, hypothetical protein |
| 3.07846 | BB2000_1467 | hypothetical protein |
| 3.18681 | BB2000_3107 | hypothetical protein |
| 3.27368 | tesB | acyl-CoA thioesterase |
| 3.3198 | BB2000_1473 | hypothetical protein |
| 3.35001 | nhaA | Na+/H+ antiporter |
| 3.38338 | BB2000_0444 | hypothetical protein |
| 3.40412 | BB2000_0426 | hypothetical protein |
| 3.49895 | pmpA | fimbrial subunit |
| 3.52689 | grcA | autonomous glycyl radical cofactor GrcA |
| 3.58086 | BB2000_1718 | hypothetical protein |
| 3.6441 | BB2000_2054 | hypothetical protein |
| 3.66735 | BB2000_2043 | hypothetical protein |
| 3.68306 | BB2000_3309 | hypothetical protein |
| 3.76035 | BB2000_0622 | hypothetical protein |
| 3.85907 | ttcA | tRNA 2-thiocytidine biosynthesis protein |
| 4.21471 | BB2000_1017 | heat shock protein |
| 4.22406 | BB2000_2810 | hypothetical protein |
| 4.60548 | BB2000_3433 | hypothetical protein |
| 4.6334 | BB2000_2357 | fimbrial operon regulator |
| 4.82102 | BB2000_1620 | hypothetical protein |
| 5.30436 | BB2000_0531 | sigma 54 modulation protein |
| 5.41104 | pheA | bifunctional chorismate mutase/prephenate dehydratase |
| 5.60442 | BB2000_0664 | hypothetical protein |
| 6.55524 | BB2000_1414 | hypothetical protein |

DRAFT

**Table S4.** Significantly differentially regulated genes between CCS01 and CCS02.

| $log_2$-fold change | BB2000 gene name | product |
|---|---|---|
| -2.87845 | BB2000_0531 | sigma 54 modulation protein |
| -2.64616 | BB2000_0669 | hypothetical protein |
| -2.63777 | BB2000_0668 | hypothetical protein |
| -2.52299 | BB2000_3110 | hypothetical protein |
| -2.43888 | BB2000_0830 | hypothetical protein |
| -2.34279 | BB2000_1017 | heat shock protein |
| -2.22256 | BB2000_0493 | phosphodiesterase |
| -2.14 | xaxB | toxin |
| -2.09976 | BB2000_2684 | chaperone protein |
| -2.05572 | BB2000_2913, BB2000_2914, BB2000_2915 | hypothetical protein, hypothetical protein, hypothetical protein |
| -2.03071 | BB2000_0667 | hypothetical protein |
| -2.02364 | BB2000_2761 | hypothetical protein |
| -2.01922 | BB2000_3468, BB2000_3469 | hypothetical protein |
| -1.9769 | BB2000_2601 | fimbrial chaperone |
| -1.87733 | BB2000_2343 | fimbrial subunit |
| -1.8413 | BB2000_0545 | hypothetical protein |
| -1.81923 | BB2000_2911 | hypothetical protein |
| -1.81253 | BB2000_1755 | hypothetical protein |
| -1.80435 | BB2000_1552 | hypothetical protein |
| -1.804 | BB2000_2920 | acyl carrier protein |
| -1.79517 | BB2000_2695 | type III secretion system protein |
| -1.79217 | BB2000_1604 | hypothetical protein |
| -1.78793 | BB2000_1182 | putative ABC transporter ATP-binding protein YbbL |
| -1.77049 | BB2000_2923 | holo-[acyl-carrier protein] synthase |
| -1.76485 | BB2000_2689 | type III secretion system protein |
| -1.76485 | BB2000_2690 | type III secretion system protein |
| -1.7577 | BB2000_2722 | microcompartments protein |
| -1.75531 | BB2000_3103 | fimbrial subunit |
| -1.73194 | rob | right origin-binding protein |
| -1.67299 | BB2000_1199 | fimbrial subunit |
| -1.66935 | BB2000_0831 | hypothetical protein |
| -1.63938 | BB2000_2697 | type III secretion system regulatory protein |
| -1.63707 | rcsA | colanic acid capsular biosynthesis activation protein (LuxR-family transcriptional regulator) |
| -1.63458 | BB2000_0445 | hypothetical protein |
| -1.6334 | BB2000_1617 | hypothetical protein |
| -1.63318 | BB2000_2355 | fimbrial subunit |
| -1.62633 | BB2000_2710 | QacE family quaternary ammonium compound efflux SMR transporter |
| -1.61553 | BB2000_2720 | microcompartments protein |
| -1.60813 | BB2000_2921 | hydroxymyristoyl-ACP dehydratase |
| -1.60662 | BB2000_2074 | hypothetical protein |
| -1.59488 | BB2000_0257 | hypothetical protein |
| -1.57819 | BB2000_2693 | type III secretion system protein |
| -1.55826 | BB2000_3389 | LysR-family transcriptional regulator |
| -1.55388 | BB2000_1393 | hypothetical protein |
| -1.55188 | BB2000_0351 | two-component sensor kinase |
| -1.54218 | BB2000_2709 | protein-tyrosine phosphatase |
| -1.54157 | terW | tellurium resistance protein |
| -1.51772 | BB2000_0408 | minor fimbrial subunit |
| -1.51663 | BB2000_0319 | hypothetical protein |
| -1.51641 | BB2000_0666 | hypothetical protein |
| -1.51027 | BB2000_2718 | hypothetical protein |
| -1.51016 | BB2000_2352 | fimbrial protein |

DRAFT

| | | |
|---|---|---|
| 1.51177 | BB2000_1438 | peptidoglycan-binding protein |
| 1.51282 | sufE | cysteine desulfuration protein |
| 1.52276 | sufS | bifunctional cysteine desulfurase/selenocysteine lyase |
| 1.52863 | sufD | cysteine desulfurase activator complex subunit SufD |
| 1.53109 | sufC | cysteine desulfurase ATPase component |
| 1.53146 | sufB | cysteine desulfurase activator complex subunit SufB |
| 1.53401 | BB2000_1822 | phage protein |
| 1.53586 | mdh | malate dehydrogenase |
| 1.53703 | hupA | DNA-binding protein HU-alpha (HU-2) |
| 1.54541 | pykA | pyruvate kinase |
| 1.5471 | mutT | nucleoside triphosphate pyrophosphohydrolase |
| 1.55277 | modB | molybdate ABC transporter permease protein |
| 1.5577 | cydX | cytochrome bd-I oxidase subunit CydX |
| 1.55954 | BB2000_0643 | hypothetical protein |
| 1.55955 | udk | uridine kinase |
| 1.56651 | sodB | superoxide dismutase [Fe] |
| 1.57226 | hupB | DNA-binding protein HU-beta |
| 1.57255 | gntY | putative DNA uptake protein |
| 1.57307 | BB2000_1358 | lipoprotein |
| 1.57311 | BB2000_0955 | transposase |
| 1.57589 | BB2000_3194 | rhodanese-like protein |
| 1.57646 | BB2000_0297 | cytoplasmic sulphur reductase |
| 1.58189 | gcp | putative DNA-binding/iron metalloprotein/AP endonuclease |
| 1.58345 | ispD | 2-C-methyl-D-erythritol 4-phosphate cytidylyltransferase |
| 1.58716 | fabI | enoyl-(acyl carrier protein) reductase |
| 1.59526 | BB2000_0840 | hypothetical protein |
| 1.5999 | cydB | cytochrome D ubiquinol oxidase subunit II |
| 1.60545 | BB2000_0949 | hypothetical protein |
| 1.61565 | zapC | type I secretion protein |
| 1.63266 | tatB | Sec-independent protein translocase |
| 1.63961 | BB2000_3319 | hypothetical protein |
| 1.64023 | pth | peptidyl-tRNA hydrolase |
| 1.65517 | tktA | transketolase |
| 1.66505 | aceF | dihydrolipoamide acetyltransferase |
| 1.66658 | aceE, pdhR | pyruvate dehydrogenase subunit E1, transcriptional regulator PdhR |
| 1.66658 | pdhR | transcriptional regulator PdhR |
| 1.66885 | secB | preprotein translocase subunit SecB |
| 1.6711 | metK | S-adenosylmethionine synthetase |
| 1.6776 | BB2000_0803 | hypothetical protein |
| 1.67923 | hfq | protein Hfq (host factor-I protein) |
| 1.68098 | hflX | putative GTPase HflX |
| 1.68212 | znuA | high-affinity zinc transporter periplasmic component |
| 1.68503 | mutL | DNA mismatch repair protein |
| 1.68523 | miaA | tRNA delta(2)-isopentenylpyrophosphate transferase |
| 1.68578 | BB2000_2664 | hypothetical protein |
| 1.69196 | BB2000_2665 | hypothetical protein |
| 1.69318 | BB2000_2194 | phosphosugar-binding regulatory protein |
| 1.69826 | dksA | DnaK transcriptional regulator DksA |
| 1.69955 | pgk | phosphoglycerate kinase |
| 1.70039 | BB2000_1964 | hypothetical protein |
| 1.70375 | pabA | para-aminobenzoate synthase component II |
| 1.70508 | BB2000_0698 | transglycosylase associated protein |
| 1.7094 | deaD | ATP-dependent RNA helicase DeaD |
| 1.7094 | BB2000_1261 | ABC transporter ATP-binding protein |
| 1.71077 | BB2000_1262 | hypothetical protein |
| 1.71077 | cspD | cold shock-like protein |

DRAFT

| | | |
|---|---|---|
| 1.71188 | rnk | nucleoside diphosphate kinase regulator |
| 1.7122 | ftsB | cell division protein FtsB |
| 1.717 | BB2000_1694 | hypothetical protein |
| 1.71725 | BB2000_1482 | hypothetical protein |
| 1.71927 | BB2000_2053 | MFS-family transporter |
| 1.72182 | BB2000_1029 | hypothetical protein |
| 1.7224 | tpiA | triosephosphate isomerase |
| 1.72559 | hpmA | hemolysin |
| 1.72932 | ipk, lolB | 4-diphosphocytidyl-2-C-methyl-D-erythritol kinase, outer membrane lipoprotein LolB |
| 1.73683 | speG | spermidine N(1)-acetyltransferase (diamine acetyltransferase) |
| 1.74322 | BB2000_1332 | translation initiation factor Sui1 |
| 1.75147 | cvpA | colicin V production protein |
| 1.75794 | atpD | F0F1 ATP synthase subunit beta |
| 1.7584 | priC | primosomal replication protein N |
| 1.75914 | BB2000_0301 | hypothetical protein |
| 1.76165 | BB2000_0592 | PhoH-like ATP-binding protein |
| 1.77641 | BB2000_1418 | transposase |
| 1.77938 | exbB | biopolymer transport protein |
| 1.78016 | exbD | biopolymer transport protein |
| 1.79239 | holE2 | DNA polymerase III, theta subunit |
| 1.79987 | rpoZ | DNA-directed RNA polymerase omega chain |
| 1.80206 | rseA | anti-RNA polymerase sigma factor SigE |
| 1.80233 | xseB | exodeoxyribonuclease VII small subunit |
| 1.81123 | BB2000_2564 | hypothetical protein |
| 1.81197 | uspA | universal stress protein A |
| 1.81253 | BB2000_1761 | methyl-accepting chemotaxis protein |
| 1.81253 | BB2000_0145 | hypothetical protein |
| 1.81257 | atpC | ATP synthase epsilon chain |
| 1.81817 | cheZ | chemotaxis regulator CheZ |
| 1.82193 | slyD | FKBP-type peptidyl-prolyl cis-trans isomerase |
| 1.82624 | apaG | ApaG |
| 1.8266 | pyrG | CTP synthetase |
| 1.82696 | glyA | serine hydroxymethyltransferase |
| 1.8298 | bfr | bacterioferritin |
| 1.83272 | BB2000_1762 | methyl-accepting chemotaxis protein |
| 1.83278 | infB | translation initiation factor IF-2 |
| 1.83343 | cstA | carbon starvation protein |
| 1.83634 | motA | flagellar motor protein MotA |
| 1.83714 | deoC | deoxyribose-phosphate aldolase |
| 1.83996 | BB2000_1050 | hypothetical protein |
| 1.84394 | ydfG | NADP-dependent L-serine/L-allo-threonine dehydrogenase |
| 1.84995 | BB2000_1308 | hypothetical protein |
| 1.85102 | BB2000_1309 | hypothetical protein |
| 1.85186 | rplQ | 50S ribosomal protein L17 |
| 1.85897 | oxaA | putative inner membrane protein translocase component YidC |
| 1.86347 | trpE | anthranilate synthase component I |
| 1.86582 | trpD | anthranilate synthase component (glutamine amidotransferase) |
| 1.8738 | trpD | anthranilate phosphoribosyltransferase |
| 1.87739 | trpC | bifunctional indole-3-glycerol phosphate synthase/phosphoribosylanthranilate isomerase |
| 1.87966 | trpB | tryptophan synthase subunit beta |
| 1.89228 | potC | spermidine/putrescine ABC transporter membrane protein |
| 1.90433 | potB | spermidine/putrescine ABC transporter membrane protein |
| 1.91043 | arnT | 4-amino-4-deoxy-L-arabinose transferase |
| 1.91522 | BB2000_0591 | putative metalloprotease |
| 1.91697 | cyoA | cytochrome o ubiquinol oxidase subunit II |

DRAFT

| | | |
|---|---|---|
| 1.91926 | BB2000_1537 | hypothetical protein |
| 1.91926 | motB | chemotaxis protein (motility protein B) |
| 1.91926 | BB2000_2177 | ArsR-family transcriptional regulator |
| 1.92053 | BB2000_1846 | hypothetical protein |
| 1.93026 | rssB | swarming motility regulation two-component system, response regulator |
| 1.93558 | accD | acetyl-CoA carboxylase subunit beta |
| 1.9375 | rplU | 50S ribosomal protein L21 |
| 1.95081 | BB2000_1827 | phage protein |
| 1.95226 | aspC | aromatic amino acid aminotransferase |
| 1.9536 | BB2000_0281 | peptidylprolyl isomerase |
| 1.95421 | chrR | chromate reductase (NADPH-dependent FMN reductase) |
| 1.95524 | lpp | major outer membrane lipoprotein (murein-lipoprotein) |
| 1.95785 | fliM | flagellar motor switch protein FliM |
| 1.98268 | fliN | flagellar motor switch protein FliN |
| 1.9859 | cfa | cyclopropane fatty acyl phospholipid synthase |
| 1.9866 | lexA | LexA repressor |
| 1.99436 | dgkA | diacylglycerol kinase |
| 1.99493 | accB | biotin carboxyl carrier protein |
| 2.00281 | BB2000_1478 | hypothetical protein |
| 2.0063 | nlpI | lipoprotein NlpI |
| 2.00797 | rpsM | 30S ribosomal protein S13 |
| 2.01102 | purA | adenylosuccinate synthetase |
| 2.01107 | mdeA | methionine gamma-lyase |
| 2.01592 | rpsU, dnaG | 30S ribosomal protein S21, DNA primase |
| 2.02025 | rpoD | RNA polymerase sigma factor RpoD |
| 2.04092 | BB2000_2511 | hypothetical protein |
| 2.04263 | eno | phosphopyruvate hydratase |
| 2.04556 | folE | GTP cyclohydrolase I |
| 2.04819 | rcsB | transcriptional regulator RcsB |
| 2.05431 | fliT | flagella protein |
| 2.05532 | BB2000_3128 | glycosyltransferase |
| 2.06105 | BB2000_0108 | cytochrome d ubiquinol oxidase subunit III |
| 2.0649 | rplM | 50S ribosomal protein L13 |
| 2.06623 | gmk | guanylate kinase |
| 2.07122 | mioC | flavodoxin |
| 2.07218 | BB2000_2866 | lipoprotein |
| 2.07572 | BB2000_3522 | hypothetical protein |
| 2.08511 | BB2000_0937 | sulphatase |
| 2.09077 | BB2000_0938 | hypothetical protein |
| 2.10985 | rrmJ | 23S rRNA methyltransferase J |
| 2.11269 | BB2000_3427 | hypothetical protein |
| 2.11325 | rpsI | 30S ribosomal protein S9 |
| 2.11884 | BB2000_3455 | hypothetical protein |
| 2.1256 | mreB | rod shape-determining protein MreB |
| 2.1256 | BB2000_2537 | phage lysis protein (holin) |
| 2.1256 | bssS | biofilm formation regulatory protein BssS |
| 2.14183 | atpF | F0F1 ATP synthase subunit B |
| 2.14525 | BB2000_3426 | hypothetical protein |
| 2.14753 | rplI | 50S ribosomal protein L9 |
| 2.14949 | BB2000_1470 | hypothetical protein |
| 2.15527 | aroK | shikimate kinase I |
| 2.16602 | groL | 60 Kda chaperonin |
| 2.17796 | fxsA | membrane protein FxsA (suppressor of F exclusion of phage T7) |
| 2.18807 | ccm | membrane protein (Ccm1 protein) |
| 2.19463 | BB2000_3025 | inorganic phosphate transporter |

DRAFT

| | | |
|---|---|---|
| 2.19755 | sthA | soluble pyridine nucleotide transhydrogenase |
| 2.19994 | grxA | glutaredoxin 1 |
| 2.20278 | tig | trigger factor |
| 2.20651 | rffH | glucose-1-phosphate thymidylyltransferase |
| 2.22152 | terZ | tellurite resistance protein |
| 2.22328 | rplO | 50S ribosomal protein L15 |
| 2.22447 | BB2000_1578 | lipoprotein |
| 2.22506 | flgL | flagellar hook-associated protein 3 (hook-filament junction protein) |
| 2.22624 | accA | acetyl-coenzyme A carboxylase carboxyl transferase subunit alpha |
| 2.22624 | thiD | phosphomethylpyrimidine kinase |
| 2.22624 | ribA | GTP cyclohydrolase II |
| 2.22686 | ompF | outer membrane porin |
| 2.22717 | gst | glutathionine S-transferase |
| 2.22872 | BB2000_1215 | PadR-family transcriptional regulator |
| 2.23943 | BB2000_1216 | hypothetical protein |
| 2.24308 | BB2000_0038 | hypothetical protein |
| 2.24596 | BB2000_0879 | hypothetical protein |
| 2.24596 | BB2000_0532 | outer membrane protein assembly complex subunit YfiO |
| 2.24816 | nusA | transcription elongation factor NusA |
| 2.24883 | BB2000_3244 | hypothetical protein |
| 2.26151 | rpsT | 30S ribosomal protein S20 |
| 2.26366 | BB2000_2949 | dihydrodipicolinate synthase-family protein |
| 2.27994 | fliS | flagellar protein FliS |
| 2.28019 | BB2000_0653 | hypothetical protein |
| 2.2813 | fkpA | FKBP-type peptidyl-prolyl cis-trans isomerase |
| 2.29202 | BB2000_3328 | hypothetical protein |
| 2.29483 | BB2000_0021 | hypothetical protein |
| 2.29493 | BB2000_1031 | lipoprotein |
| 2.29551 | trxA | thioredoxin |
| 2.29561 | BB2000_0725 | probable transporter |
| 2.30025 | BB2000_1041 | hypothetical protein |
| 2.30546 | BB2000_0946 | hypothetical protein |
| 2.31165 | cmk | cytidylate kinase |
| 2.31834 | ptsI | phosphoenolpyruvate-protein phosphotransferase |
| 2.32395 | BB2000_1825 | phage protein |
| 2.33486 | BB2000_2653 | chitin binding protein |
| 2.33739 | rpmB | 50S ribosomal protein L28 |
| 2.34384 | msrB | peptide methionine sulfoxide reductase |
| 2.34565 | BB2000_3399 | hypothetical protein |
| 2.35404 | rplA | 50S ribosomal protein L1 |
| 2.35487 | rmf | ribosome modulation factor |
| 2.35971 | lrp | leucine-responsive transcriptional regulator |
| 2.36114 | ftsK | cell division protein (DNA translocase) |
| 2.37187 | BB2000_0771 | recombination factor protein RarA |
| 2.38002 | uspE | universal stress protein UspE |
| 2.38596 | fnr | fumarate/nitrate reduction transcriptional regulator |
| 2.3922 | rpsR | 30S ribosomal protein S18 |
| 2.40355 | BB2000_0573 | hypothetical protein |
| 2.40947 | BB2000_0792 | hypothetical protein |
| 2.41343 | BB2000_1844 | lipoprotein |
| 2.41575 | secG | protein-export membrane protein |
| 2.43869 | rpsO | 30S ribosomal protein S15 |
| 2.44193 | BB2000_1419 | hypothetical protein |
| 2.45316 | BB2000_1829 | phage antitermination protein |
| 2.48477 | BB2000_1828 | phage holin (lysis protein) |

DRAFT

| | | |
|---|---|---|
| 2.48477 | sodA | superoxide dismutase [Mn] |
| 2.48802 | csaA | protein secretion chaperone |
| 2.49849 | BB2000_1070 | transcriptional regulator |
| 2.50114 | BB2000_1071 | fimbrial subunit |
| 2.50209 | thrS | threonyl-tRNA synthetase |
| 2.50447 | infC | translation initiation factor IF-3 |
| 2.50743 | rplT | 50S ribosomal protein L20 |
| 2.5128 | pheS | phenylalanyl-tRNA synthetase alpha chain |
| 2.51809 | pheT | phenylalanyl-tRNA synthetase subunit beta |
| 2.52214 | ihfA | integration host factor subunit alpha |
| 2.53924 | btuD | vitamin B12 import ATP-binding protein |
| 2.55988 | arnB | UDP-4-amino-4-deoxy-L-arabinose–oxoglutarate aminotransferase |
| 2.56167 | arnC | undecaprenyl phosphate 4-deoxy-4-formamido-L-arabinose transferase |
| 2.56238 | arnA | bifunctional UDP-glucuronic acid decarboxylase/UDP-4-amino-4-deoxy-L-arabinose formyltransferase |
| 2.56558 | BB2000_1083 | polysaccharide deacetylase |
| 2.57237 | arnT | 4-amino-4-deoxy-L-arabinose transferase |
| 2.58146 | BB2000_1085 | hypothetical protein |
| 2.58165 | BB2000_1086 | hypothetical protein |
| 2.58412 | gltA | type II citrate synthase |
| 2.58699 | trmD | tRNA (guanine-N1)-methyltransferase |
| 2.5922 | rpsD | 30S ribosomal protein S4 |
| 2.59794 | fliF | flagellar MS-ring protein |
| 2.6035 | fliH | flagellar assembly protein H |
| 2.60543 | fliI | flagellum-specific ATP synthase |
| 2.61728 | BB2000_1824 | phage protein |
| 2.61805 | ompA | outer membrane protein A |
| 2.62741 | priB | primosomal replication protein N |
| 2.63659 | dps | DNA starvation/stationary phase protection protein Dps |
| 2.63831 | metJ | transcriptional repressor protein MetJ |
| 2.64642 | atpI | F0F1 ATP synthase subunit I |
| 2.65607 | holD | DNA polymerase III subunit psi |
| 2.66553 | flgD | basal-body rod modification protein |
| 2.66643 | fis | DNA-binding protein Fis |
| 2.66717 | dusB | tRNA-dihydrouridine synthase B |
| 2.66717 | BB2000_2967 | iron ABC transporter, substrate-binding protein |
| 2.66717 | BB2000_3143 | hypothetical protein |
| 2.66717 | BB2000_2855 | signal sensing protein |
| 2.67152 | nusB | transcription antitermination protein NusB |
| 2.67459 | flgE | flagellar hook protein FlgE |
| 2.68283 | BB2000_1204 | hypothetical protein |
| 2.68443 | BB2000_1756 | hypothetical protein |
| 2.68991 | acpP | acyl carrier protein |
| 2.6928 | fabF | 3-oxoacyl-(acyl carrier protein) synthase II |
| 2.69332 | ppsA | phosphoenolpyruvate synthase |
| 2.70672 | infA | translation initiation factor IF-1 |
| 2.70948 | aat | leucyl/phenylalanyl-tRNA–protein transferase |
| 2.71304 | cydC | cysteine/glutathione ABC transporter membrane/ATP-binding component |
| 2.72113 | cydD | cysteine/glutathione ABC transporter membrane/ATP-binding component |
| 2.73101 | atpB | ATP synthase A chain |
| 2.73299 | BB2000_2806 | intracellular sulfur oxidation protein |
| 2.73299 | BB2000_2950 | hypothetical protein |
| 2.73444 | tufB | elongation factor Tu |

DRAFT

| | | |
|---|---|---|
| 2.73554 | BB2000_3500 | acetyltransferase |
| 2.73716 | crp | cAMP-regulatory protein |
| 2.75141 | ydgA | hypothetical protein |
| 2.75171 | suhB | inositol monophosphatase |
| 2.77913 | rpmF | 50S ribosomal protein L32 |
| 2.80008 | ribH | 6, 7-dimethyl-8-ribityllumazine synthase |
| 2.82143 | BB2000_2828 | hypothetical protein |
| 2.83558 | atpE | F0F1 ATP synthase subunit C |
| 2.84894 | crr | glucose-specific PTS system component |
| 2.85653 | proQ | putative solute/DNA competence effector |
| 2.86937 | rpsC | 30S ribosomal protein S3 |
| 2.87388 | ihfB | integration host factor subunit beta |
| 2.87807 | ndk | nucleoside diphosphate kinase |
| 2.88299 | icd | isocitrate dehydrogenase |
| 2.89118 | lpxC | UDP-3-O-[3-hydroxymyristoyl] N-acetylglucosamine deacetylase |
| 2.89813 | rplR | 50S ribosomal protein L18 |
| 2.90167 | rplY | 50S ribosomal protein L25 |
| 2.90167 | BB2000_1795 | hypothetical protein |
| 2.90167 | flgA | flagella basal body P-ring formation protein |
| 2.90167 | rnpA | ribonuclease P |
| 2.90167 | rpsH | 30S ribosomal protein S8 |
| 2.90167 | fusA | elongation factor G (EF-G) |
| 2.9107 | rimM | 16S rRNA-processing protein RimM |
| 2.9165 | tolB | translocation protein TolB |
| 2.95158 | rplN | 50S ribosomal protein L14 |
| 2.95832 | budA | alpha-acetolactate decarboxylase |
| 2.95947 | rpsP | 30S ribosomal protein S16 |
| 2.97887 | BB2000_3459 | hypothetical protein |
| 2.98135 | prsA | ribose-phosphate pyrophosphokinase |
| 2.98876 | rpmD | 50S ribosomal protein L30 |
| 3.01302 | rplE | 50S ribosomal protein L5 |
| 3.01933 | BB2000_2388 | oxidoreductase |
| 3.01933 | ribB | 3, 4-dihydroxy-2-butanone 4-phosphate synthase |
| 3.03474 | rplP | 50S ribosomal protein L16 |
| 3.05733 | rpsQ | 30S ribosomal protein S17 |
| 3.06819 | rpsG | 30S ribosomal protein S7 |
| 3.07376 | mipA | MltA-interacting protein precursor |
| 3.07972 | dadB | alanine racemase, catabolic |
| 3.08307 | dadA | D-amino acid dehydrogenase small subunit |
| 3.08856 | pal | peptidoglycan-associated outer membrane lipoprotein |
| 3.09235 | BB2000_1584 | transcriptional regulator |
| 3.09882 | flgB | flagellar basal body rod protein FlgB |
| 3.11079 | BB2000_0342 | transcriptional regulator |
| 3.11822 | rpsJ | 30S ribosomal protein S10 |
| 3.12247 | fliD | flagellar capping protein |
| 3.14106 | sdhA | succinate dehydrogenase flavoprotein subunit |
| 3.14414 | sucA | 2-oxoglutarate dehydrogenase E1 component |
| 3.15071 | sucB | dihydrolipoamide succinyltransferase component of 2-oxoglutarate dehydrogenase complex |
| 3.16157 | sucC | succinyl-CoA synthetase subunit beta |
| 3.20086 | sucD | succinyl-CoA synthetase alpha chain |
| 3.20903 | BB2000_0639 | hypothetical protein |
| 3.21287 | BB2000_2819 | methyl-accepting chemotaxis protein |
| 3.21951 | rplV | 50S ribosomal protein L22 |
| 3.24112 | BB2000_1483 | hypothetical protein |
| 3.26729 | BB2000_1956 | lipoprotein |

DRAFT

| | | |
|---|---|---|
| 3.26729 | BB2000_2387 | hypothetical protein |
| 3.26729 | rplX | 50S ribosomal protein L24 |
| 3.31065 | BB2000_0110 | hypothetical protein |
| 3.3325 | BB2000_2655 | hypothetical protein |
| 3.3325 | rplD, rplW, rplB | 50S ribosomal protein L4, 50S ribosomal protein L23, 50S ribosomal protein L2 |
| 3.35328 | fliA | flagellar biosynthesis sigma factor |
| 3.38664 | rplC | 50S ribosomal protein L3 |
| 3.39462 | BB2000_1381 | outer membrane protein (attachment invasion locus protein) |
| 3.42274 | emrR | transcriptional repressor MprA |
| 3.43981 | tufB | elongation factor Tu |
| 3.46147 | fliE | flagellar hook-basal body complex protein |
| 3.50226 | rpsN | 30S ribosomal protein S14 |
| 3.5091 | rpmH | 50S ribosomal protein L34 |
| 3.51141 | fumC | fumarate hydratase |
| 3.51141 | BB2000_1316 | hypothetical protein |
| 3.51141 | intB | prophage integrase |
| 3.51141 | flgM, flgN | anti-sigma28 factor FlgM, flagella synthesis protein |
| 3.51141 | cspA | cold shock protein |
| 3.51141 | BB2000_1717 | hypothetical protein |
| 3.51141 | fliZ | flagella biosynthesis protein FliZ |
| 3.51141 | BB2000_1015 | lipase |
| 3.51782 | BB2000_0873 | hypothetical protein |
| 3.5361 | BB2000_0874 | hydrolase |
| 3.54258 | cspB | cold shock protein |
| 3.56173 | ddg | cold-induced palmitoleoyl transferase |
| 3.56379 | BB2000_0877 | hypothetical protein |
| 3.59905 | BB2000_0878 | hypothetical protein |
| 3.59909 | BB2000_1586 | hypothetical protein |
| 3.6236 | idrA | IdrA |
| 3.66151 | BB2000_1097, BB2000_1098, BB2000_1099 | hypothetical protein, fimbrial protein, Fimbrial usher protein |
| 3.70279 | BB2000_1100 | fimbrial chaperone |
| 3.72567 | BB2000_1102 | fimbrial subunit |
| 3.74534 | BB2000_1103 | fimbrial protein |
| 3.75525 | BB2000_1104 | fimbrial protein |
| 3.76036 | BB2000_3499 | lipoprotein |
| 3.79038 | BB2000_0148 | hypothetical protein |
| 3.8123 | BB2000_0579 | hypothetical protein |
| 3.83155 | BB2000_0942 | hypothetical protein |
| 4.02273 | BB2000_2810 | hypothetical protein |
| 4.04203 | BB2000_0184 | hypothetical protein |
| 4.04203 | BB2000_0509 | hypothetical protein |
| 4.10138 | BB2000_1721 | hypothetical protein |
| 4.10384 | BB2000_2492 | hypothetical protein |
| 4.27611 | BB2000_2619 | hypothetical protein |
| 4.31151 | BB2000_2639 | hypothetical protein |
| 4.35192 | BB2000_2663 | hypothetical protein |
| 4.37037 | BB2000_3387 | hypothetical protein |
| 4.37723 | BB2000_0126 | hypothetical protein |
| 4.50389 | BB2000_0881 | hypothetical protein |
| 4.50389 | BB2000_1054 | hypothetical protein |
| 4.50389 | BB2000_1488 | hypothetical protein |
| 4.50389 | BB2000_1610 | hypothetical protein |
| 4.50389 | BB2000_1770 | hypothetical protein |
| 4.50389 | BB2000_2320 | hypothetical protein |
| 4.5988 | BB2000_3091 | hypothetical protein |

DRAFT
